# A quartz crystal microbalance method to quantify the size of hyaluronan and other glycosaminoglycans on surfaces

**DOI:** 10.1101/2022.02.28.482386

**Authors:** Sumitra Srimasorn, Luke Souter, Dixy E. Green, Lynda Djerbal, Ashleigh Goodenough, James A. Duncan, Abigail R. E. Roberts, Xiaoli Zhang, Delphine Débarre, Paul L. DeAngelis, Jessica C. F. Kwok, Ralf P. Richter

**Author notes:** These authors contributed equally (Sumitra Srimasorn, Luke Souter).

## Abstract

Hyaluronan (HA) is a major component of peri- and extra-cellular matrices and plays important roles in many biological processes such as cell adhesion, proliferation and migration. The abundance, size distribution and presentation of HA dictate its biological effects and are also useful indicators of pathologies and disease progression. Methods to assess the molecular mass of free-floating HA and other glycosaminoglycans (GAGs) are well established. In many biological and technological settings, however, GAGs are displayed on surfaces, and methods to obtain the size of surface-attached GAGs are lacking. Here, we present a method to size HA that is end-attached to surfaces. The method is based on the quartz crystal microbalance with dissipation monitoring (QCM-D) and exploits that the softness and thickness of films of grafted HA increase with HA size. These two quantities are sensitively reflected by the ratio of the dissipation shift (Δ*D*) and the negative frequency shift (-Δ*f*) measured by QCM-D upon the formation of HA films. Using a series of size-defined HA preparations, ranging in size from ∼2 kDa tetrasaccharides to ∼1 MDa polysaccharides, we establish a monotonic yet non-linear standard curve of the Δ*D*/-Δ*f* ratio as a function of HA size, which reflects the distinct conformations adopted by grafted HA chains depending on their size and surface coverage. We demonstrate that the standard curve can be used to determine the mean size of HA, as well as other GAGs, such as chondroitin sulfate and heparan sulfate, of preparations of previously unknown size in the range from 1 to 500 kDa, with a resolution of better than 10%. For polydisperse samples, our analysis shows that the process of surface-grafting preferentially selects smaller GAG chains, and thus reduces the average size of GAGs that are immobilised on surfaces comparative to the original solution sample. Our results establish a quantitative method to size HA and other GAGs grafted on surfaces, and also highlight the importance of sizing GAGs directly on surfaces. The method should be useful for the development and quality control of GAG-based surface coatings in a wide range of research areas, from molecular interaction analysis to biomaterials coatings.

## 1. Introduction

Polysaccharides of the glycosaminoglycan (GAG) family are major constituents of extracellular matrices. The presence of these complex carbohydrates in virtually all vertebrate tissues implicates their diverse functions and importance: GAGs regulate many different processes in tissue physiology and pathology, such as in ovulation and fertilisation for mammalian reproduction [1], in tissue development [2], in the regulation of neuronal plasticity [3], in tissue mechanical functions (*e*.*g*., in cartilage [4, 5]), in infection, inflammation and immunity [6-8], and in cancer biology [9].

The GAG family has five members: hyaluronan (HA; the only GAG that is constitutively non-sulfated), heparan sulfate (HS; including the highly sulfated form and drug heparin), chondroitin sulfate (CS), dermatan sulfate (DS) and keratan sulfate (KS). All GAGs are polymers of unbranched polysaccharides composed of repeating disaccharides, negatively charged at physiological pH (with sulphated GAGs having the highest charge density of all biomacromolecules known), well soluble in water and mechanically compliant. Owing to these common physical properties, GAG chains are generally thought to intrinsically lack a defined secondary or higher order structure, but on their own rather dynamically sample a wide range of low energy conformations [3], and upon binding to various proteins may adopt more distinct conformations [7, 10, 11].

Hence, in contrast to folded proteins the diversity of GAG functions is not encoded in its higher order structure but rather in (i) the chemical nature and varying sulfation of the comprising disaccharides of the GAG chain (as these dictate which extracellular proteins they bind), (ii) the core proteins to which sulfated GAGs are typically covalently tethered (thus forming proteoglycans), (iii) the number of GAG chains attached to the core proteins (thus controlling their hydrodynamic properties) and (iv) GAG chain size (molecular mass). The mass of HA, for example, can vary over several orders of magnitude, from a few kDa for the smallest oligosaccharides to many MDa for the largest polysaccharides. The variability in size arises from distinct HA synthases (HAS1, HAS2 and HAS3) which produce HA of different size distributions, and from hyaluronidases or reactive oxygen species (*e*.*g*., in inflammation) which degrade HA to different levels dependent on tissue context. The chain length of HA appears to influence cell apoptosis, proliferation and mobility [12, 13]. During inflammation, HA of high molecular mass is generally considered anti-inflammatory but low molecular mass pro-inflammatory. Similarly, proteoglycans and their sulfated GAG chains vary in size depending on tissue context and function.

In light of the wide range of GAG sizes and their biological importance, the ability to quantify the size of GAGs is key to progress in GAG biology and the use of GAGs in medical, pharmaceutical and cosmetic applications. Several methods are now well established to quantify the mass of HA and other GAGs in the solution phase [14]. Multi-angle light scattering (MALS) when coupled with size exclusion chromatography (SEC) or field flow fractionation (FFF) provides weight-averaged (*M*_W_) and number-averaged (*M*_n_) molecular masses, and thus also the polydispersity index (PDI = *M*_W_ / *M*_n_) of a sample without the need for a size standard but requires relatively large amounts of sample (typically tens to hundreds of μg for SEC-MALS [14]). Gas-phase electrophoretic mobility molecular analysis (GEMMA) provides mean size and size dispersity, and requires a much smaller amount of sample (pg range) [15]. Gel electrophoresis (GE) can be performed in any biochemistry lab and can be adapted to virtually any GAG size (with sample needs in the upper ng to lower μg range, dependent on size distribution and staining method), but requires a set of size standards for analysis [14, 16, 17]. Importantly, GEMMA and GE are strongly affected by GAG charge in addition to size which limits the use of these methods for sulphated GAG polysaccharides with varying charge distribution. More recently, solid-state nanopore sensors have emerged for the quantification of HA mass [18]. Analysing one molecule at a time, the technique provides detailed size distribution information and requires relatively little sample (ng amounts).

All of the above techniques provide mass information for molecules that are free in the solution phase. In many biological settings, however, GAGs are displayed on surfaces or other scaffolding structures, forming films of varying thickness. For example, HA may be retained on the cell surface through one end (*e*.*g*., by HA synthases) or through multiple attachment points along the chain contour (*e*.*g*., by HA receptors such as CD44 or LYVE-1 [19, 20]), and sulfated GAGs such as CS, DS and HS are typically tethered *via* one end to their core protein [3]. ‘End-on’ or ‘side-on’ attachments of GAGs are also being widely used for biomedical applications such as implant coatings [21-23], biomaterial scaffolds [24, 25] and nanoparticles for theranostics [26-29], and for biophysical assays to study protein and virus binding to GAGs and GAG cross-linking by proteins [11, 30-33]. In these applications, GAG attachment is typically *via* one end to recapitulate its native, flexible conformation and because this enables the GAG surface density to be easily tuned.

There is currently no well-established method for size analysis of end-attached HA and other GAGs. Atomic force microscopy can trace the contour of individual GAG chains and resolve the size of the GAG populations. This technique can be applied to individual GAG chains [34], or to CSPGs where GAG chains are attached to proteoglycan core proteins [35], when firmly attached along their entire contour to a surface. However, applications of this approach are rather limited owing to the need for sample drying and a rather poor control over GAG deposition. HA and other GAGs may be stripped off surfaces and scaffolds for subsequent solution phase size analysis [36]. However, on-surface analysis of GAG size would not require stripping and concentration steps necessary for solution-phase analysis, thus simplifying analysis and avoiding potential artefacts (such as accidental GAG chain degradation), and also enabling in-line quality control of GAG-coated surfaces prior to their further use.

We here present a method to quantify the mean size of GAGs that are attached with one end to a planar surface. The method is based on quartz crystal microbalance with dissipation monitoring (QCM-D), a technique that is widely used for the analysis of macromolecular interactions and thin films at surfaces [37-39]. Through measuring changes in the resonance frequency (Δ*f*) and the energy dissipation (Δ*D*; the rate of decay of the induced oscillations) of a shear-oscillating quartz crystal sensor with sub-second time resolution, QCM-D can monitor changes in the mass per unit surface area (areal mass density) indicating binding, along with the mechanical characteristics of the surface-bound film in real time. The mechanical properties ultimately reflect on the conformation and flexibility of surface-bound molecules and/or the molecular interactions within the surface-attached film.

To a first approximation, in QCM-D an increased areal mass density produces a negative Δ*f*, and an increased ‘softness’ produces a positive Δ*D*. The areal mass density includes any solvent that is hydrodynamically coupled to the surface-adhered film in the sensor’s shear oscillation (which in amount can far exceed the water or ion molecules chemically associated with the macromolecular film [38, 40]), and for sufficiently dense or laterally homogeneous films QCM-D thus reports the film thickness [37]. Δ*D* is though also affected by the areal mass density, and interface mechanical properties also impact on Δ*f* for soft and thick films. This implies that in the general case of soft films Δ*f* and Δ*D* responses need to be considered jointly to quantify film thickness and mechanical properties [37]. Numerical analysis of Δ*f* and Δ*D* values measured at several overtones (*i*.*e*., harmonics of the sensor’s resonance) with viscoelastic models can indeed provide quantification of film thickness and viscoelasticity, but this procedure is not appropriate for monolayers of discrete molecules (owing to additional energy dissipation pathways not accounted for by the models [37, 41]). Moreover, viscoelastic modelling also has limitations for laterally homogeneous films: careful fitting is required to ascertain meaningful numbers with their confidence intervals are extracted from the data [37, 42], and for very soft films (including GAG films) it is not generally possible to determine film thickness and viscoelastic properties simultaneously from QCM-D data with tight confidence intervals owing to too strong parameter correlation.

To overcome these limitations, Du and Johannsmann introduced the Δ*D*/-Δ*f* ratio as a quantitative measure of the elastic compliance (a measure of softness) for films that are ultrathin (*i*.*e*., in the range of a few nm) yet laterally homogeneous [43, 44]. For thicker films, the proportionality is lost but the Δ*D*/−Δ*f* ratio remains a useful measure to quantify film softness [37, 42, 43]. Compared to viscoelastic models, a key benefit of the Δ*D*/-Δ*f* ratio is that it can be computed easily and transparently from the original QCM-D data. The Δ*D*/-Δ*f* ratio can also provide information about monolayers of discrete molecules, on flexible linker regions between the molecules and the surface, or within the molecule [37, 41]. Interestingly, Gizeli *et al*. demonstrated that the Δ*D*/-Δ*f* ratio parallels changes in the intrinsic viscosity, a measure of molecular shape in the solution phase, of surface-tethered DNA chains, and that the Δ*D*/-Δ*f* ratio can discriminate one-end tethered nucleic acid (DNA) polymer chains by their size [43, 45, 46].

Considering that GAGs and nucleic acid polymers are both flexible, linear and well solvated in aqueous solution, we rationalised that QCM-D should be similarly sensitive to the size of GAGs. Indeed, we here show that the sensitivity of QCM-D for mechanical properties enables QCM-D to discriminate GAGs by their size. We establish the Δ*D*/-Δ*f* ratio as a measure for GAG size and develop a simple and robust method to determine the mean size of one-end tethered GAG chains based on the Δ*D*/-Δ*f* ratio using a few μg or less of material with an analysis time in the range of minutes.

## 2. Materials and methods

### 2.1 Materials

Lyophilized 1,2-dioleoyl-sn-glycero-3-phosphocholine (DOPC) and 1,2-dioleoyl-sn-glycero-3-phosphoethanolamine-N-(Cap Biotinyl) (DOPE-cap-B) were purchased from Avanti Polar Lipids (Alabaster, USA). Lyophilised streptavidin (SAv) was purchased from Sigma-Aldrich.

Quasi-monodisperse size-defined hyaluronan, either plain (HA) or with a biotin tag at the reducing end (HA-b), was supplied as lyophilized powder from Hyalose (Oklahoma City, USA) or custom made as described previously [47]. Quasi-monodisperse size-defined heparosan (Hep-b) and chondroitin (C0-b) with a biotin tag at their reducing ends were custom made for this study using synchronized, stoichiometrically controlled chemoenzymatic reactions. For HEP-b, a heparosan trisaccharide amine derivative was extended with recombinant MBP-PmHS1 synthase using UDP-GlcNAc and UDP-GlcA (Sigma) [48]. For C0-b, a HA tetrasaccharide amine [47] was extended with recombinant PmCS1-704 synthase [49]) using UDP-GalNAc and UDP-GlcA [50, 51]. These amine-tagged GAGs were biotinylated with biotin-LC-sulfoNHS reagent (Thermo) at 30× molar excess in 50 mM HEPES, pH 7.2 overnight. The target GAG-biotin polymer was then purified by ethanol precipitation (2.5 volumes, 0.1 M NaCl final concentration) and repeated ultrafiltration against water in a 3 kDa MWCO spin-unit (Millipore). Size-defined heparan sulfate (HS-b) oligosaccharides with a biotin tag at their reducing end were custom made as described previously [52]. See **Table 1** for details and references regarding size-defined GAGs.

**Table 1.**
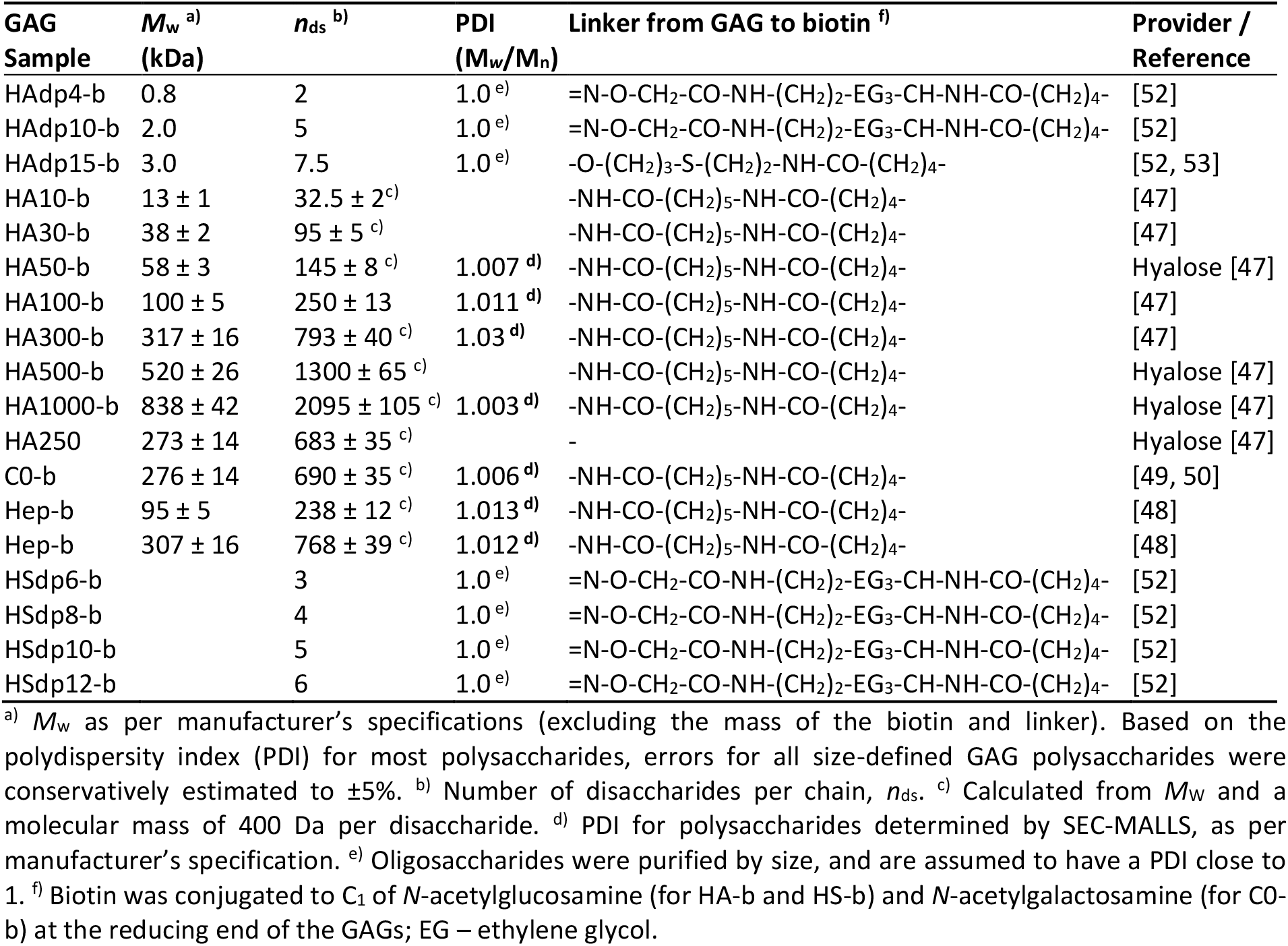
Size-defined GAG samples used in this study.

Preparations of chondroitin sulfate (CS), including with varying sulfation levels (CS-A, CS-C, CS-D and CS-E), dermatan sulfate (DS) and heparan sulfate (HS) (all extracted and purified from animal tissues), and HA (purified from microbial fermentation and size fractionated), were purchased from commercial providers; see **Table 2** for details. These preparations, which are known to show a rather large size distribution, were biotinylated at their reducing end by oxime ligation; see Supplementary Methods for details.

**Table 2.**
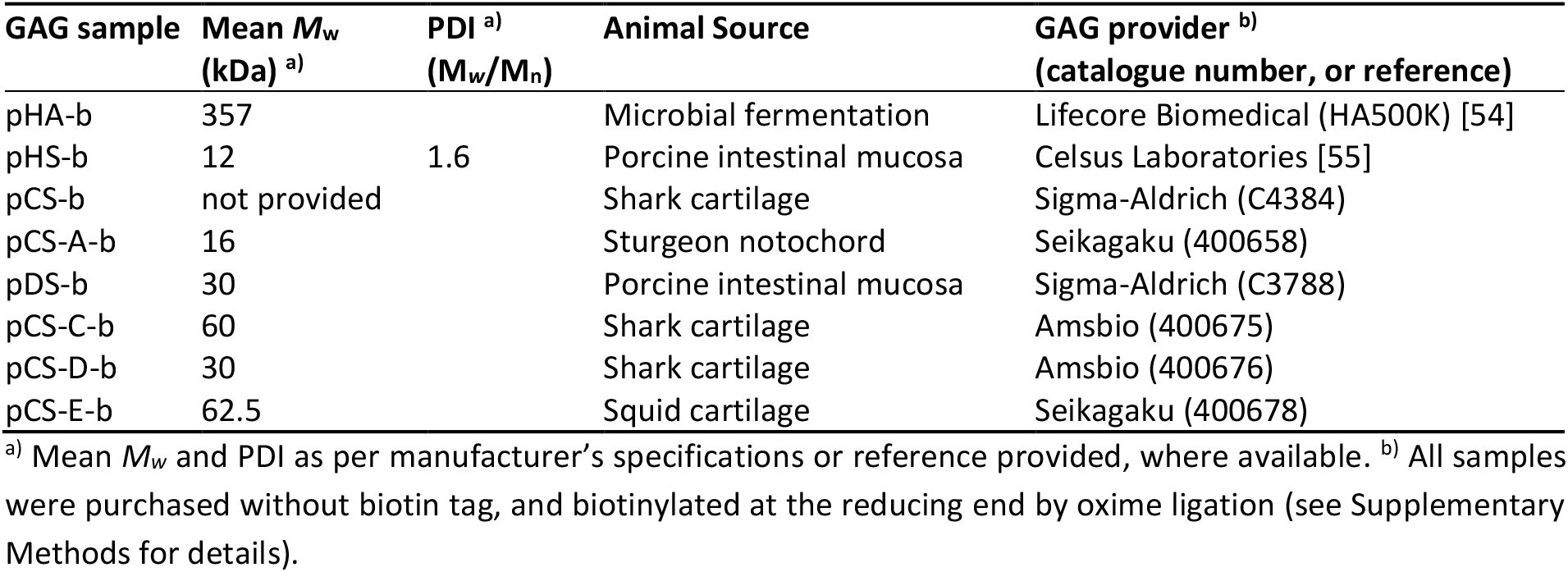
Polydisperse GAG polysaccharide samples used in this study.

### 2.2 Sample preparation

Working buffer for all QCM-D experiments consisted of 10 mM HEPES, pH 7.4, 150 mM NaCl (*i*.*e*., physiological salt levels and pH) prepared in ultrapure water (uH_2_O; resistivity 18.2 MΩ·cm^-1^).

Small unilamellar vesicles displaying biotin (b-SUVs) were prepared as previously described [56] with modifications. Briefly, lipids in chloroform were mixed at a ratio of 95 mol-% DOPC and 5 mol-% DOPE-cap-B at a total amount of 5 µmol, and dried under a stream of nitrogen gas followed by drying in a vacuum desiccator for 2 h. The lipid mixture was re-suspended in working buffer at 2 mg·mL^-1^, and homogenised by five cycles of freezing, thawing and vortexing. To obtain SUVs, the lipid suspension was subjected to tip sonication in pulse mode (1 s on / 1 s off) for 30 min with refrigeration. The SUV suspension was then centrifuged at 12,100 × *g* for 10 min to remove titanium debris (shed from the sonicator tip), and stored at 4°C under nitrogen gas until use.

Lyophilized SAv was dissolved in uH_2_O at 1 mg·mL^-1^ and stored at -20 °C until use. GAGs were re-suspended in uH_2_O at stock concentrations between 0.1 and 1 mg·mL^-1^. GAGs of M_W_ < 250 kDa were vortexed for rapid sample homogenisation; larger GAGs were not vortexed but incubated at 4 °C for 2 h without shaking to avoid chain fragmentation due to mechanical shear. All GAG samples were stored at -20 °C until use.

### 2.3 Quartz crystal balance with dissipation monitoring (QCM-D)

QCM-D measurements on silica-coated sensors (QSX303; Biolin Scientific, Västra Frölunda, Sweden) were performed with a Q-Sense E4 system (Biolin Scientific) equipped with 4 independent flow modules, connected to a syringe pump (Legato; World Precision Instruments, Stevenage, UK) to deliver a fluid flow of 10 to 20 µL·min^-1^. The working temperature was set to 24 °C. Changes in resonance frequency (Δ*f*_*i*_) and dissipation (Δ*D*_*i*_) were acquired from six overtones (*i* = 3, 5, 7, 9, 11 and 13, corresponding to resonance frequencies of *f*_*i*_ ≈ 5, 15, 25, 35, 45, 55 and 65 MHz). Results from overtone *i* = 3 are presented unless stated otherwise, and frequency shifts are presented normalised by the overtone number (Δ*f* = Δ*f*_*i*_/*i*). All other overtones provided qualitatively similar data.

### 2.4 Polyacrylamide gel electrophoresis (PAGE)

GAG samples were analysed by PAGE for independent crude confirmation of their size and size distribution. Each GAG sample of molecular mass > 250 kDa (0.125 µg for size-defined samples, 3 µg for polydisperse samples) was mixed with FACE loading agent (a mixture of glycerol and uH_2_O at a ratio of 1:3 *v*/*v*, and 1 mg·mL^-1^ bromophenol blue) and 1× Tris-borate buffer (0.05 M Trizma base, 0.06 M boric acid, pH 8.3) at volumes of 10 µL, 3 µL and 2 µL, respectively. GAG polysaccharides of lower molecular mass (1 µg for size-defined samples, 3 µg for polydisperse samples) were mixed with a different FACE loading agent (a mixture of glycerol, DMSO and uH_2_O at a ratio of 1:2:7 *v*/*v*, and 1 mg·mL^-1^ bromophenol blue) and 2× Tris-borate buffer at volumes of 10 µL, 3 µL and 2 µL, respectively. Samples and appropriate molecular mass markers (HA ladder; Hyalose) were loaded onto a Tris-borate gel and run on 2× Tris-borate buffer at 300 V and 4 °C. Gels were fixed with 10% ethanol in 1× Tris-borate buffer for 30 min prior to staining with Alcian Blue solution (0.4% *w*/*v* Alcian Blue 8GX (Sigma Aldrich), 40% *v*/*v* methanol and 8% *v/v* acetic acid in uH_2_O) for visualization of GAGs. De-staining was performed with an aqueous solution containing 40% *v*/*v* methanol and 8% *v*/*v* acetic acid.

## 3. Results

### 3.1 GAG brushes provide defined anchorage, suitable for GAG sizing on surfaces

Here, we introduce QCM-D as a method to assess the size of GAG chains that are anchored with one end to a planar surface. GAGs with a biotin tag at the reducing end (GAG-b) were anchored to a streptavidin (SAv)-coated supported lipid bilayer (SLB). The anchorage approach illustrated in **Figure 1a** (left) is well established [57, 58], and provides for site-specific attachment (*i*.*e*., ‘grafting’ *via* biotin) without secondary interactions of the GAG chains with the SLB/SAv-coated surface.

**Figure 1.**
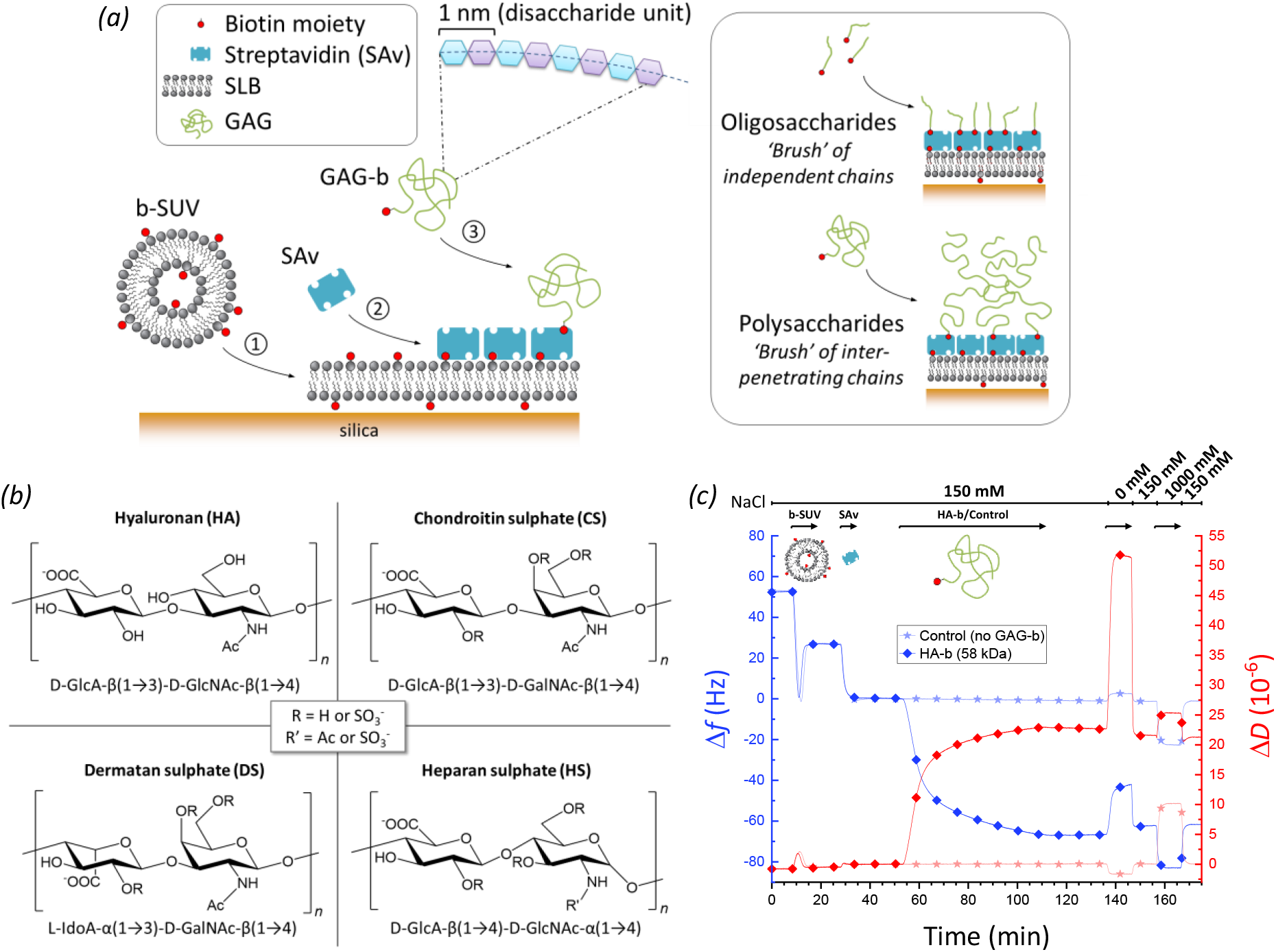
Experimental setup for GAG-sizing by QCM-D. *(a)* Schematic displaying the surface functionalisation steps: ➀ Formation of a supported lipid bilayer (SLB) on the quartz crystal sensor surface (silica) from small unilamellar vesicles containing 5 mol-% biotinylated lipids (b-SUVs); ➁ Formation of a streptavidin (SAv) monolayer (reference surface); ➂ Anchorage of GAGs of various sizes *via* a biotin at their reducing end (GAG-b). All molecules are drawn roughly to scale, except the GAG polysaccharides for which the linear dimensions can exceed SAv by 10-fold and more. *(b)* Disaccharide unit structures of the GAG types studied, including positions of potential sulfations (for CS, DS and HS; additionally, GlcA units of HS may be epimerised into IdoA). *(c)* Representative QCM-D experiment monitoring all typical incubation steps. Shifts in frequency (Δ*f*; *blue lines*) and dissipation (Δ*D*; *red lines*) for the third overtone (*i* = 3) are shown for an experiment with HA-b (58 kDa; *lines with diamonds*) and for an experiment without any GAG-b (control; *lines with stars*). The start and the duration of sample incubation steps are indicated by the *arrows* above the graph along with the NaCl concentration step profile. Incubation conditions: buffer – 10 mM HEPES, pH 7.4, with NaCl as indicated; b-SUVs – 50 μg/mL; SAv – 20 μg/mL; HA-b (58 kDa) - 5 μg/mL.

The conformation of the GAG chains on the sensor depends on the GAG size and surface coverage, and also on the ion concentration (*vide infra*). GAG oligosaccharides (≲20 monosaccharide units) have a contour length (1 nm per disaccharide in the chain) comparable in magnitude to the persistence length (a measure of chain flexibility; 4.1 nm for HA [59], and likely similar for other GAGs, at physiological salt concentration and pH) and therefore will present extended yet dynamically bending conformations in solution. Surface grafting is not expected to substantially affect the conformation of these chains, *i*.*e*., they will adopt a range of random orientations, and only slightly interact with each other even at the highest attainable surface densities (**Figure 1a**, upper right). On the other hand, GAG polysaccharides (≳20 monosaccharide units) have a much larger contour length and thus form extended ‘random coils’ in solution. When anchored to the surface at low coverage, the random coil conformation is largely preserved, but chains will interpenetrate and repel each other (due to the negative charges on the numerous GlcA monosaccharides) at sufficiently high anchorage density, thus inducing preferential chain stretching away from the surface (**Figure 1a**, lower right) [60-62]. We here refer to films of grafted and stretched oligosaccharide/polysaccharide chains as GAG ‘brushes’, consistent with established terminology for grafted polymers [63].

We monitored all surface functionalisation steps and the brush formation process for various GAG types and sizes (**Figure 1b, Table 1** and **Table 2**) using QCM-D (**Figure 1c**) at close-to-physiological pH (7.4) and ionic strength (150 mM NaCl), to ascertain the surface preparation proceeded correctly and to quantify the QCM-D responses (frequency shifts, Δ*f*, and dissipation shifts, Δ*D*) upon GAG-b binding. For any of the incubation steps, a decrease in Δ*f* (**Figure 1c**, blue lines) indicates an increase in the sensed mass on the surface (which includes hydrodynamically coupled solvent) while the associated Δ*D* changes (**Figure 1c**, red lines) reflect on the softness of the biomolecular films [57, 64]. The QCM-D responses evidenced the formation of a proper SLB (**Figure 1c**, 7 – 17 min) and a densely packed monolayer of SAv (**Figure 1c**, 27 – 33 min) as a robust platform for GAG brush formation (see refs. [65, 66] and Supplementary Methods for detailed analyses of these processes). As a whole, the SAv-on-SLB film generates very little dissipation shift, *i*.*e*., it is essentially sensed as a rigid film by QCM-D. Subsequent incubation with GAGs with biotin tag consistently led to an additional decrease in frequency and a marked increase in dissipation, as exemplified in **Figure 1c** (lines with diamonds, 53 – 111 min) for HA-b with a molecular mass of 58 kDa. These shifts were absent in a control experiment utilising plain, unmodified HA (273 kDa) lacking a biotin tag (**Figure S1**, lines with stars), demonstrating that GAGs attach to the surface exclusively *via* specific biotin-SAv interactions. Moreover, the GAG brushes were stable over time, as seen by the absence of further frequency and dissipation shifts upon rinsing in working buffer (**Figure 1c**; 111 – 137 min).

### 3.2 Size-defined HAs reveal how steric hindrance of GAG brushes influences the magnitude of QCM-D responses

To establish how sensitive QCM-D is to GAG size and therefore suitable for sizing GAG chains, we compared a set of 10 HA-b samples with well-defined sizes ranging from 2 disaccharides (*n*_ds_ = 2) to 838 kDa (*n*_ds_ ≈ 2100; **Table 1**). The full set of QCM-D data are shown in **Figure S2**. We chose HA because of its perfectly regular disaccharide structure (with no additional modification) and because size-defined samples of HA were available over a wide range of chain sizes. Polyacrylamide gel electrophoresis showed all quasi-monodisperse polysaccharide samples as defined and narrow bands (**Figure S3**), indicating no or very little contamination with low molecular mass residues that could have resulted from fragmentation during handling.

We first inspected the QCM-D responses at maximal HA-b binding (*i*.*e*., at the end of the HA incubation process; **Figure 2**). Interestingly, both Δ*f*_min_ nor Δ*D*_max_ varied non-monotonically with HA size: the magnitudes of Δ*f*_min_ and Δ*D*_max_ were largest at an intermediate size (Δ*f*_min_ = -72 Hz for HA-b 38 kDa, and Δ*D*_max_ = 23 × 10^−6^ for HA-b 58 kDa, respectively), but decreased towards smaller and larger sizes. The size-dependent trends observed can be attributed to two opposing effects of GAG size. On the one hand, the QCM-D response per chain is expected to increase with size owing to the increased mass (and hydrodynamically coupled solvent) of larger chains, at comparable grafting densities. On the other hand, the chain size negatively impacts the grafting density that can be attained in the experiment; the GAG chains on forming a brush impose a steric barrier to the access of additional polysaccharide chains to the surface, and thus gradually reduce the binding rate as the brush becomes denser [57, 67]. This barrier effect is expected to be negligible for very short oligosaccharide chains yet very pronounced for long polysaccharide chains due to the larger hydrodynamic volume that they occupy and the larger steric hindrance they exert.

**Figure 2.**
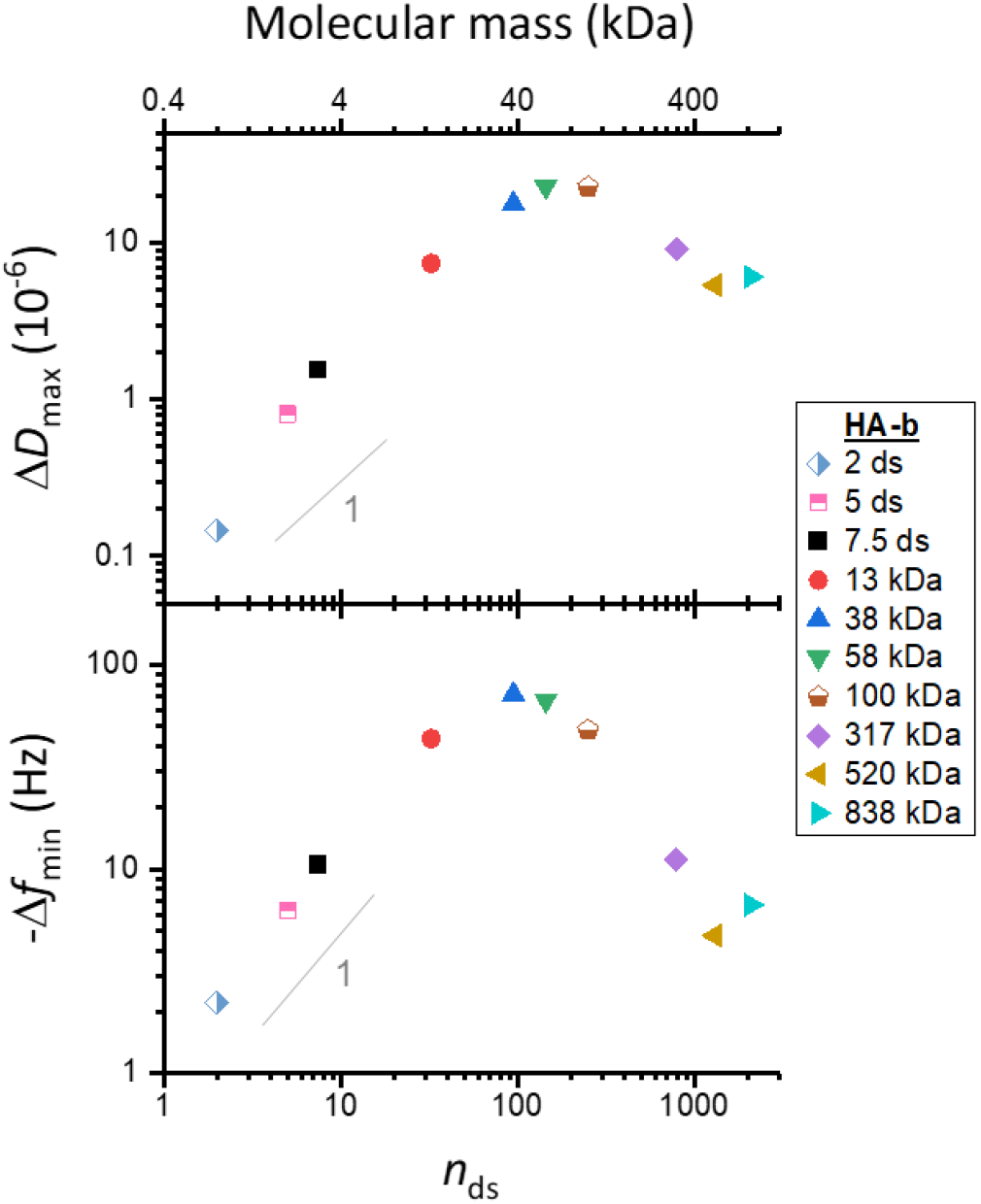
Maximal QCM-D responses vary non-monotonically with HA size. Double-logarithmic plots with shifts in dissipation (Δ*D*_max_) and frequency (-Δ*f*_min_; *i* = 3) at maximum GAG binding as a function of the mean number of disaccharides, *n*_ds_, per chain. Estimated experimental uncertainties (accounting for baseline drift upon HA binding: 0.67 × 10^−6^ for Δ*D* and 0.33 Hz for Δ*f* per hour) are smaller than the symbol size and not shown. Grey lines with a slope of one are shown for reference: -Δ*f*_min_ increases roughly linearly with HA size for small HA sizes. Incubation conditions: HA-b oligosaccharides (2, 5 and 7.5 ds; all chains ≲3 kDa) and HA-b polysaccharides (13, 38, 58 and 317 kDa) – 5 μg/mL; HA-b 100 kDa – 10 µg/mL; HA-b 520 kDa and HA-b 838 kDa – 20 μg/mL; see **Figure S2** for original data.

Our data (**Figure S2**) are entirely consistent with longer saccharides exerting steric hindrance, as evidenced by reduced binding rates and lower grafting densities with increasing HA size. HA-b binding saturated within 10 min or less for all HA-b oligosaccharides (*n*_ds_ = 2, 5 and 7.5) and also for the smallest HA-b polysaccharide (13 kDa, *n*_ds_ ≈ 33) tested, indicating that these short chains readily occupy all available biotin binding sites on the SAv monolayer. The binding site density for a similarly prepared surface was previously shown to be 7.8 pmol/cm^2^ [53], corresponding to a root-mean-square (rms) distance between GAG anchors of *d*_rms_ = 4.6 nm. In this regime, where the films are dense and thus relatively rigid, -Δ*f*_min_ increases roughly linearly with HA size (**Figure 2**). In contrast, the binding rate of all other longer HA-b chains gradually decreased with coverage and did not saturate during the incubation times selected in our experiments (40 to 60 min). For the largest HA-b chains tested (317, 520 and 838 kDa), the barrier to binding increased to such an extent that binding virtually stalled after about 30 min of HA-b incubation. For HA-b (838 kDa), for example, we have previously reported rms distances between anchor points of *d*_rms_ > 50 nm under conditions similar to the ones used here [68, 69], implying that molar anchorage densities are more than 100-fold reduced for the largest GAG chains compared to oligosaccharides. The reduced grafting densities also render the less dense brushes formed with the largest HA-b chains softer (*vide infra*), and this characteristic further attenuates the QCM-D frequency shift.

In conclusion, the reduction in grafting density outweighs the mass effect of chain size on the QCM-D response. Moreover, the lack of monotonic trends makes the Δ*f* or Δ*D* values alone unsuitable for establishing a clear relationship (*i*.*e*., a standard curve) between GAG size and QCM-D response.

### 3.3. The ΔD/-Δf ratio enables robust HA sizing

In light of the complexity in the dependence of HA-b size on the frequency and dissipation shifts when each parameter is analysed on their own, we instead explored the Δ*D*/-Δ*f* ratio. The Δ*D*/-Δ*f* ratio is a measure of the elastic compliance (or ‘softness’) for ultrathin yet homogenous films [43, 44]. In an alternative approach, Gizeli *et al*. demonstrated that the Δ*D*/-Δ*f* ratio parallels changes in the intrinsic viscosity (a measure of molecular shape in the solution phase) of surface-grafted DNA chains, and that the Δ*D*/-Δ*f* ratio is approximately independent of surface coverage [43, 45, 46]. Building on these findings, we hypothesised that this parameter could provide a simple approach to measure GAG size.

Using the QCM-D responses for each HA-b size, the Δ*D*/-Δ*f* ratio was calculated and plotted as a function of the negative frequency shift (-Δ*f*) which here serves as a proxy for relative surface coverage (**Figure 3**). A clear trend can be discerned, with the Δ*D*/-Δ*f* ratio showing a pronounced increase as a function of HA size (for any given –Δ*f* value). We also noticed changes in the Δ*D*/-Δ*f* ratio with surface coverage (*i*.*e*., as a function of -Δ*f*, for any given HA size), although these were typically less pronounced than the dependence on size: for HA oligosaccharides and polysaccharides of intermediate sizes (≤ 58 kDa), only a weak monotonic decrease in Δ*D*/-Δ*f* with -Δ*f* was generally noted. For HA chains of higher molecular weight (> 58 kDa), coverage effects were more pronounced, with an initial increase preceding the phase of decreasing Δ*D*/-Δ*f*. Notably, the Δ*D*/-Δ*f* ratios for HA of 317, 520 and 838 kDa were comparable at the highest surface coverages but clearly distinct at low coverages. Thus, the sensitivity of the Δ*D*/-Δ*f* ratio to HA size at low coverage provides a better distinction for sizing GAGs.

**Figure 3.**
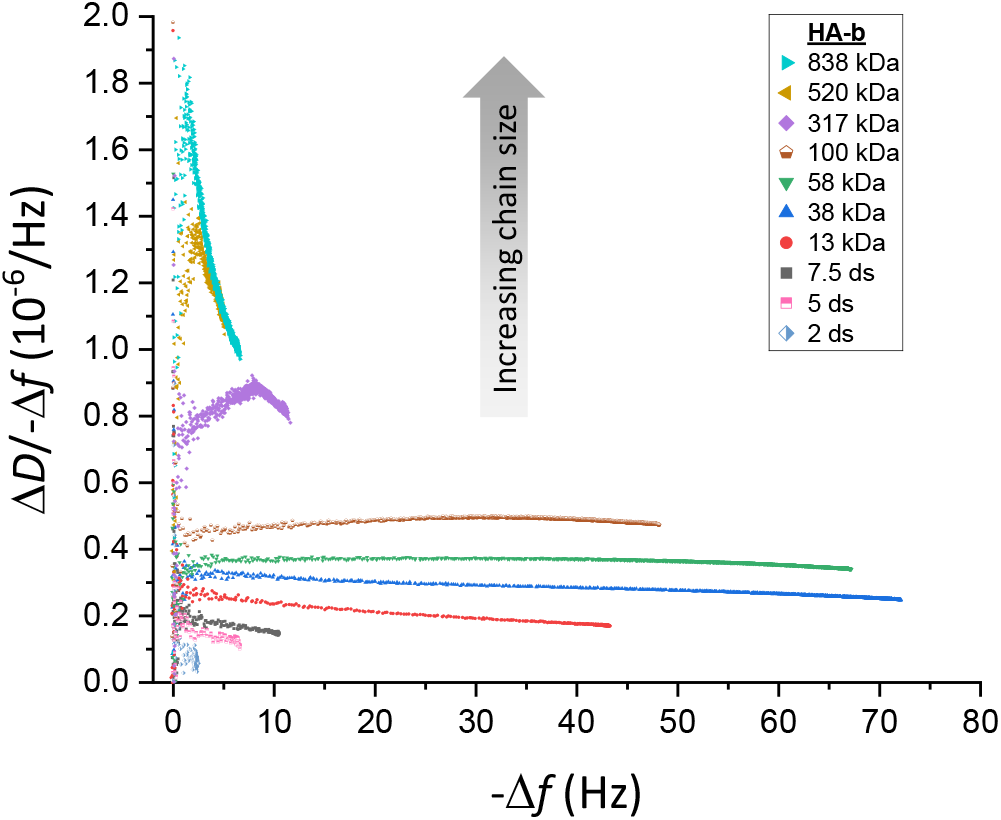
Δ*D*/-Δ*f* analysis robustly discriminates HA sizes. Parametric plots of Δ*D*/-Δ*f vs*. –Δ*f* (*i* = 3) during the formation of brushes of HA with defined sizes (as indicated with symbol and colour code; see **Figure S2** for original data.

### 3.4. Establishing a standard curve for sizing HA

Having demonstrated that the Δ*D*/-Δ*f* ratio is a sensitive predictor for HA size over a wide size range, we aimed to establish a standard curve for practical use in GAG sizing applications. Using Δ*D*/-Δ*f* ratios at maximum polysaccharide coverage for size analysis is suboptimal, because the coverage varies between experiments with incubation conditions and these coverage variations impact on the Δ*D*/-Δ*f* ratio (**Figure 3**). Hence, a specific target frequency for GAG sizing would be preferable.

We chose -Δ*f* = 2.5 Hz for Δ*D*/-Δ*f* ratio determination, which provides several benefits. First, the choice of a specific frequency eliminates the issue of coverage dependence in the Δ*D*/-Δ*f* ratio, and thus enhances the robustness of the data. Second, the Δ*D*/-Δ*f* ratio at -Δ*f* = 2.5 Hz is accessible for all GAG sizes, from the shortest oligosaccharides to the longest polysaccharides. Third, it is a good balance that combines an acceptable signal-to-noise ratio in the Δ*D*/-Δ*f* values (noise becomes excessive as -Δ*f* approaches zero, as seen in **Figure 3**) with a minimal amount of sample and/or experimental time (which increase towards higher coverage, or -Δ*f* values).

Figure 4. shows the Δ*D*/-Δ*f* ratio (at -Δ*f* = 2.5 Hz) as a function of HA size. The mean and the standard error of the mean (*blue spheres* with error bars) were here determined from between two and four independent experiments per HA size. The errors are generally small, indicating good reproducibility of the Δ*D*/-Δ*f* ratio (see **Figure S4** for results of all individual experiments). HA size is expressed in *n*_ds_.

**Figure 4.**
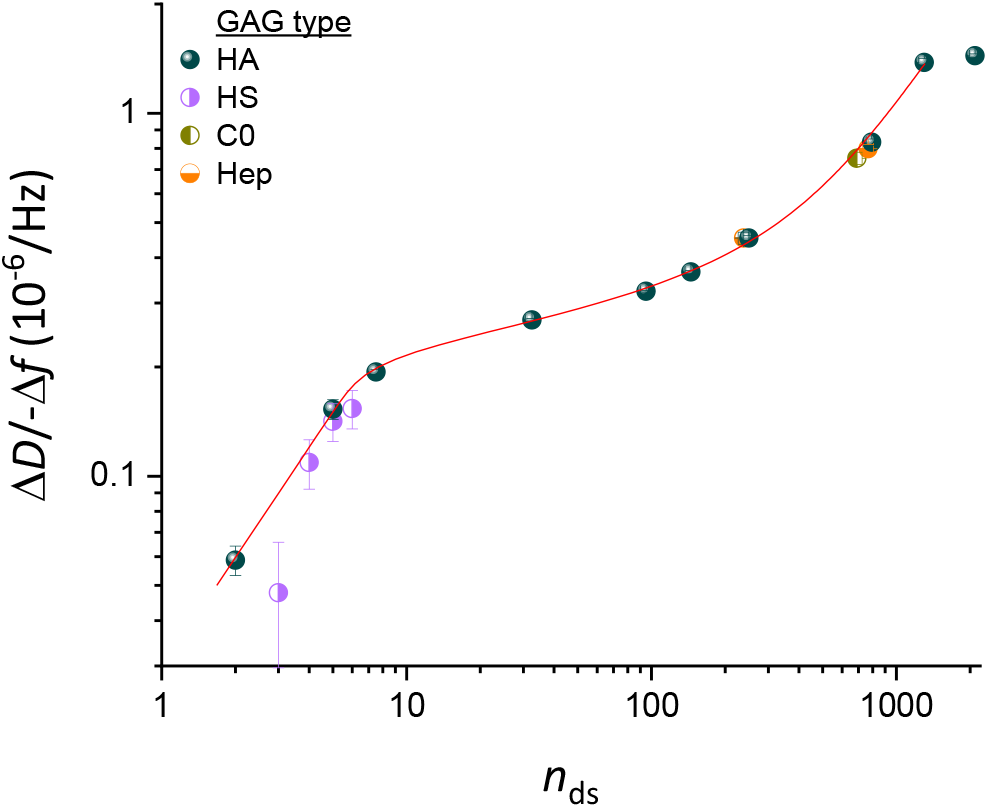
Standard curve for sizing HA, and extension to other GAG types. Double logarithmic plot of Δ*D*/-Δ*f* (*i* = 3) at low GAG surface density (-Δ*f* = 2.5 Hz) as a function of the mean number of disaccharides per GAG-b chain, *n*_ds_ (mean ± 5%, except for oligosaccharides which were taken to be pure in size, see **Table 1**). Data for all size-defined HA-b samples (*blue spheres*) represent mean ± standard error of the mean from between two and four independent experiments (see **Figure S4** for details). The data for HA up to *n*_ds_ = 1300 are faithfully reproduced and interpolated by **Equation 1** (*red line*), representing the standard curve for GAG sizing. Data for size-defined preparations of other GAG types (*half-filled circles*; see **Figure S6** for original data) are also seen to map onto the standard curve: four heparan sulfate (HS-b; *violet*) oligosaccharides, two heparosan (Hep-b; *orange*) or one chondroitin (C0-b; *dark yellow*) polysaccharide. Symbols and error bars for non-HA GAGs represent individual experiments with the mean ± standard deviation time-averaged over -Δ*f* values ranging between 2 and 3 Hz.

As expected, a clear trend of Δ*D*/-Δ*f* ratios monotonically increasing with HA size can be discerned. However, the Δ*D*/-Δ*f* ratio dependence is rather complex: a sharp increase for oligosaccharides (*n*_ds_ = 2 to 7.5) is followed by a shallower slope for small and intermediate-sized polysaccharides (up to *n*_ds_ = 800), a renewed sharp increase between *n*_ds_ = 800 and 1300, and a plateau is effectively attained for the largest polysaccharides (*n*_ds_ > 1300). The red line in **Figure 4** corresponds to the function

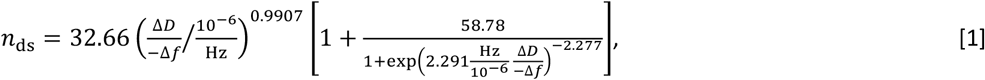

where the five numerical values were determined through fitting to the data for all HA sizes up to *n*_ds_ = 1300 (520 kDa). The choice of this function was empirical (see Section 4.3 for a discussion of the physical origin of the curve shape) yet it reproduces the experimental data faithfully and thus provides a useful tool for data interpolation. Note that **Equation 1** expresses *n*ds as a function of Δ*D*/-Δ*f*, rather than *vice versa*, to facilitate the determination of the GAG size from an experimentally measured Δ*D*/-Δ*f* value. The average (*i*.*e*., root-mean-square) deviation in *n*_ds_ between the experimental data and the interpolating fit (**Figure S5**) was below 2%, and the maximal deviation was below 9%, across the HA size range considered, indicating that **Equation 1** faithfully reproduces the experimental data and thus can serve as an accurate standard curve.

Figure 4. was deliberately drawn as a double-logarithmic plot. In this presentation, the local slopes of the standard curve represent the rate of relative change in Δ*D*/-Δ*f* (d(Δ*D*/−Δ*f*)/(Δ*D*/−Δ*f*) = ln(Δ*D*/−Δ*f*)) as a function of relative change in HA size (d*n*ds/*n*ds = In *n*ds), and are an effective measure of size sensitivity of the Δ*D*/-Δ*f* ratio. The slope *α* = d ln(Δ*D*/−Δ*f*)/d In *n*_ds_ is largest (α ≈ 1.0) for oligosaccharides up to *n*_ds_ = 10 and for polysaccharides around *n*_ds_ ≈ 10^3^, and the method therefore is most sensitive in these size ranges. For intermediate sizes, the size sensitivity is somewhat reduced (down to α ≈ 0.2 for *n*_ds_ ≈ 30), and above *n*_ds_ ≈ 1300 (or *M*_W_ ≈ 500 kDa) it is virtually lost (α ≈ 0).

The size resolution of our standard curve can be estimated from the slopes *α* and the resolution of the Δ*D*/-Δ*f* ratio, which is represented by the standard deviation across multiple experiments. Analysis of the data indicates a resolution of better than 10% up to *n*_ds_ ≈ 1300, with the exception of the smallest HA oligosaccharide (*n*_ds_ = 2) where the resolution is 17% (**Figure S5**).

Our QCM-D setup monitors changes in resonance frequency and dissipation at higher overtones (*i* ≥ 5), in addition to *i* = 3. Comparative analysis (**Figure S5** shows results for *i* = 3, 5 and 7) revealed that the size sensitivity of Δ*D*/-Δ*f* was broadly similar, albeit reduced for the largest HA sizes, for higher overtones. Together, the results demonstrate that Δ*D*/-Δ*f* values at -Δ*f* = 2.5 Hz for *i* = 3 provide a robust and effective discrimination for HA sizes up to ∼500 kDa.

### 3.5. The standard curve for sizing HA can also be used for other GAG types

The standard curve in **Figure 4** (red line) was established exclusively with size-defined HA. Is this curve also representative for other GAG types? All other GAG types share similar basic disaccharide subunits (*i*.*e*., a uronic acid linked to a hexosamine) and hydrodynamic properties with HA to merit a direct comparison. To address this question, we probed size-defined preparations of other GAG types.

As other examples of GAG polysaccharides, we used heparosan (Hep-b 100 kDa and 307 kDa) and chondroitin (C0-b 276 kDa) (see **Figure S6** for time-resolved QCM-D data and parametric plots of Δ*D*/-Δ*f vs*. -Δ*f*). These regular polysaccharides recapitulate the basic monosaccharide sequences of heparan sulfates/heparin (except for a potential epimerisation of GlcA into IdoA; **Figure 1b**) and chondroitin sulfates, respectively, but are the unsulfated biosynthetic precursors. These carbohydrates thus possess a similar linear structure and the same charge density as HA (*i*.*e*., one charge per disaccharide), which makes them ideally suited to probe the impact of potential variations of the polysaccharide ‘backbone’, such as intrinsic chain flexibility, on the Δ*D*/-Δ*f* ratio. **Figure 4** shows that the Δ*D*/-Δ*f* ratios (at -Δ*f* = 2.5 Hz) for Hep-b (half-filled orange circles at *n*_ds_ = 238 and 768, respectively) and C0-b (half-filled dark yellow circle at *n*_ds_ = 690) are close to the values for HA-b of similar size, indicating that the standard curve of the defined quasi-monodisperse HA provides a suitable proxy to size non-sulfated GAG polysaccharides other than HA.

As oligosaccharides, we used heparan sulfate preparations of 3, 4, 5 or 6 disaccharides (see **Figure S6** for time-resolved QCM-D data and parametric plots). These oligosaccharides were isolated to monodispersity from an enzymatic digest of heparan sulfate with an average of 1.4 sulfates per disaccharide unit [55], yet are variable in their level and pattern of sulfation. **Figure 4** shows that the Δ*D*/-Δ*f* ratios for most of the HS-b oligosaccharides (half-filled violet circles) are close to the HA-b standard curve. The Δ*D*/-Δ*f* ratio for the smallest HS oligosaccharide was slightly lower than the interpolated HA data, but overall it appears that the relation of the Δ*D*/-Δ*f* ratio to size (at 150 mM NaCl) is not substantially impacted by sulfation for oligosaccharides. Taken together, these findings suggest that the standard curve established with HA can be applied to determine the size of non-sulfated GAGs of any type, and also of sulfated GAG polysaccharides.

### 3.6. Charge-mediated repulsion between GAG chains also affects the QCM-D response, but this effect is small at 150 mM NaCl

For polysaccharides, it was impossible to directly probe the effect of sulfation on the Δ*D*/-Δ*f* ratio because size-defined sulfated GAG polysaccharides are not available. Recognising that the main effect of sulfation on GAG morphology is due to an increased charge, we explored varying the NaCl concentration in our experiments as an indirect way to assess the effect of GAG sulfation on the Δ*D*/-Δ*f* ratio. All GAGs are negatively charged at neutral pH owing to the carboxyl groups (one per each disaccharide for all GAGs), and for sulfated GAGs the sulfation contributes a substantial number of additional chargeable groups: up to three per each disaccharide, depending on GAG type and degree of sulfation (**Figure 1b**). Whilst added charges on the GAGs are expected to increase the repulsion between GAG chains, this effect can be counteracted by increasing the ionic strength of the solution. To test how GAG charge impacts the sensing of GAG brushes by QCM-D, we thus probed responses of brushes made from size-defined HA-b to low (0 mM) and high (1000 mM) NaCl concentrations whilst maintaining the pH at 7.4 with 10 mM HEPES. Whilst we expect the repulsion between GAG chains to be strongly enhanced at 0 mM NaCl, all charge effects on GAG morphology are effectively eliminated at 1 M NaCl [64, 70].

These tests were performed at the end of each experiment following HA-b brush formation in 150 mM NaCl (see **Figure 1c**, >130 min, for an example, and **Figure S2** for a complete set of QCM-D data covering all HA sizes). A set of control experiments were required to ascertain that the HA films retain their brush morphology and integrity upon transient exposure to high and low salt conditions. Experiments with non-biotinylated HA (273 kDa; **Figure S1**) confirmed that the SAv-on-SLB surface remained inert to HA binding and that HA-b was surface-bound exclusively *via* the reducing-end biotin tag, under all tested salt conditions. QCM-D signals also fully recovered upon return to 150 mM NaCl for most HA brushes (**Figure S2**). Notable exceptions were dense brushes made of intermediate HA sizes (*i*.*e*., 13 and 38 kDa, and to a minor extent also 58 kDa), where low salt induced so much swelling and pressure within the brush that it promoted the release of a fraction of the anchored HA chains (see **Figure S7** for a detailed analysis of this effect). When comparing data over a range of salt concentrations, we ascertained that QCM-D responses to salt changes were not affected by changes in HA grafting density. Further control experiments were conducted on bare SAv-on-SLB surfaces (**Figure 1c** and **Figure S2**, lines with stars) to quantify the effect of salt changes on the QCM-D response that are not related to HA but arise from the sensitivity of the QCM-D to changes in solution viscosity and density upon variations of the NaCl concentration. These effects from changes in solution properties were subtracted from the QCM-D responses obtained with HA brushes to calculate the net effect due to the impact of ionic strength variations on the HA brushes (see Supplementary Methods for details).

Figure 5. compares the net Δ*D*/-Δ*f* ratios for HA brushes for 0 mM, 150 mM or 1000 mM NaCl across all HA sizes. Note that we here consider the Δ*D*/-Δ*f* ratios at maximal surface coverage; it would be challenging to test the effect of salt at low surface coverage (-Δ*f* between 2 to 3 Hz, as established in **Figure 4**) because of excessively large experimental errors associated with the buffer exchanges relative to the QCM-D response for HA binding. Clear differences are noticeable when comparing the data for 0 mM to 1000 mM NaCl, demonstrating that the Δ*D*/-Δ*f* ratio in general is sensitive to GAG charge in addition to GAG size. Effects are most obvious for the HA polysaccharides, where the Δ*D*/-Δ*f* ratio increases with decreasing ionic strength, likely due to brush swelling (see **Figure S2**, legend, for a related analysis of the complex trends observed for Δ*D* and Δ*f* individually). This tendency appears to reverse for GAG oligosaccharides, although the effect is relatively small and it is unclear if it is significant considering the experimental errors.

**Figure 5.**
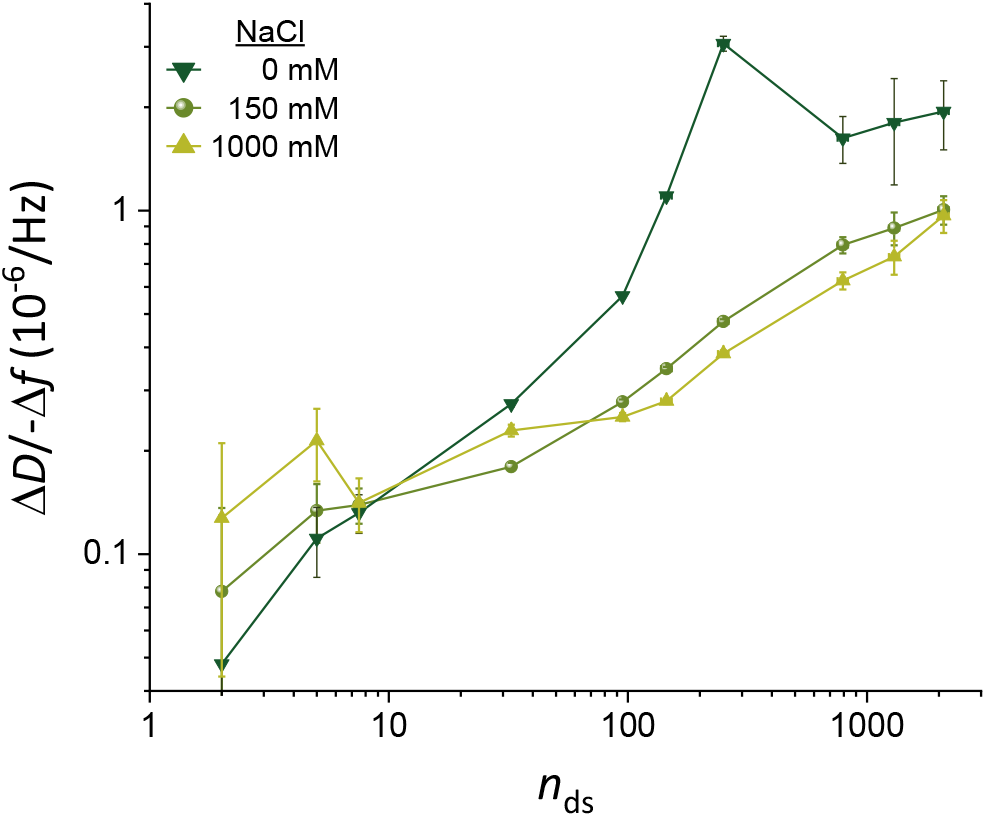
Effect of ionic strength on Δ*D*/-Δ*f* ratios. Double logarithmic plot of Δ*D*/-Δ*f* (*i* = 3) as a function of the mean number of disaccharides (*n*_ds_) per HA chain, for NaCl concentrations of 0 mM, 150 mM and 1000 mM (as indicated with symbol and colour code). Representative data for one experiment per HA size are shown. The mean values represent the Δ*D*/-Δ*f* ratio measured at high GAG surface densities (*i*.*e*., the maximal surface densities that were attained in the experiments and stable at 0 mM NaCl; see **Figure S2** for original data). The error bars account for baseline drifts during GAG binding. In the cases of 0 and 1000 mM NaCl, solution effects on the QCM-D response (*i*.*e*., owing to the effect of salt on solution density and/or viscosity) needed to be corrected for with the aid of control data, and uncertainties associated with this correction are also included in the error bars (see **Figure 1c** and Supplementary Methods for details).

The Δ*D*/-Δ*f* ratios at 150 mM, on the other hand, were comparable to 1000 mM NaCl within experimental error for small HA chains (≤38 kDa) as well as for the largest HA chains (≥520 kDa), and only moderately increased (by up to 27%) for intermediate HA sizes (58, 100 and 317 kDa). The relatively small differences demonstrate that charge effects are largely screened at 150 mM NaCl and that Δ*D*/-Δ*f* values at 150 mM NaCl primarily report on GAG size, although some caution is advised for GAG sizes in the range of many tens of kDa to a few hundred kDa. This finding provides further support to the use of 150 mM NaCl (as already used in Figure 4) for GAG sizing applications, including for sulfated GAG polysaccharides.

### 3.7. Application examples for GAG sizing

We present a few simple examples to illustrate the benefits of the developed method to determine the mean molecular mass of surface-grafted GAGs. In most cases of practical relevance, GAG samples are derived from natural sources and purified to varying degrees. With the exception of oligosaccharides, purified GAGs from natural sources (*e*.*g*., animal tissues, cells and bacteria) retain a substantial degree of polydispersity (illustrated in **Figure S8**) even with the most advanced current size fractionation methods. Thus, we explored if the mean size of surface-grafted GAGs faithfully recapitulate the size distribution of the original GAG solution from which a GAG brush is being formed.

We tested this with a solution of polydisperse HA with a mean molecular mass of 357 kDa according to the manufacturer (**Table 2**). Gel electrophoresis confirmed the broad dispersity of this HA reagent (**Figure S8a** and **Figure 6b**) after biotinylation at the reducing end, although the most abundant size appeared to be somewhat higher (∼500 kDa); half-maximal staining intensities above background were reached at approximately 200 kDa on the lower end, and >800 kDa at the higher end (the gel did not allow reliable determination of the upper limit). In contrast, the Δ*D*/-Δ*f* ratio obtained with this sample (**Figure 6a**; see **Figure S9a** for the time-resolved QCM-D data) provided an effective HA mean size on the surface of 150±40 kDa (**Figure 6b**), through comparison with the standard curve (**Figure 4** and **Equation 1**). This example illustrates that the surface bound technology can unintentionally favour the binding of smaller GAGs in a polydisperse sample leading to the mean GAG size on the surface being substantially skewed lower than in the starting solution. One likely reason for preferential binding of smaller GAG molecules with lower hydrodynamic radius is their faster diffusion to the surface. In addition, the size skewing will be exacerbated by the barrier properties of the forming GAG brush; the barrier will be most pronounced for the longest chains in the GAG pool and the brush thus effectively sieves out shorter chains for surface binding. The latter effect should be particularly pronounced for large GAGs (≥100 kDa; see Section 4.3 for a detailed discussion), and we propose that this mechanism explains the more than two-fold reduction in effective mean size upon surface anchorage for the large polydisperse HA.

**Figure 6.**
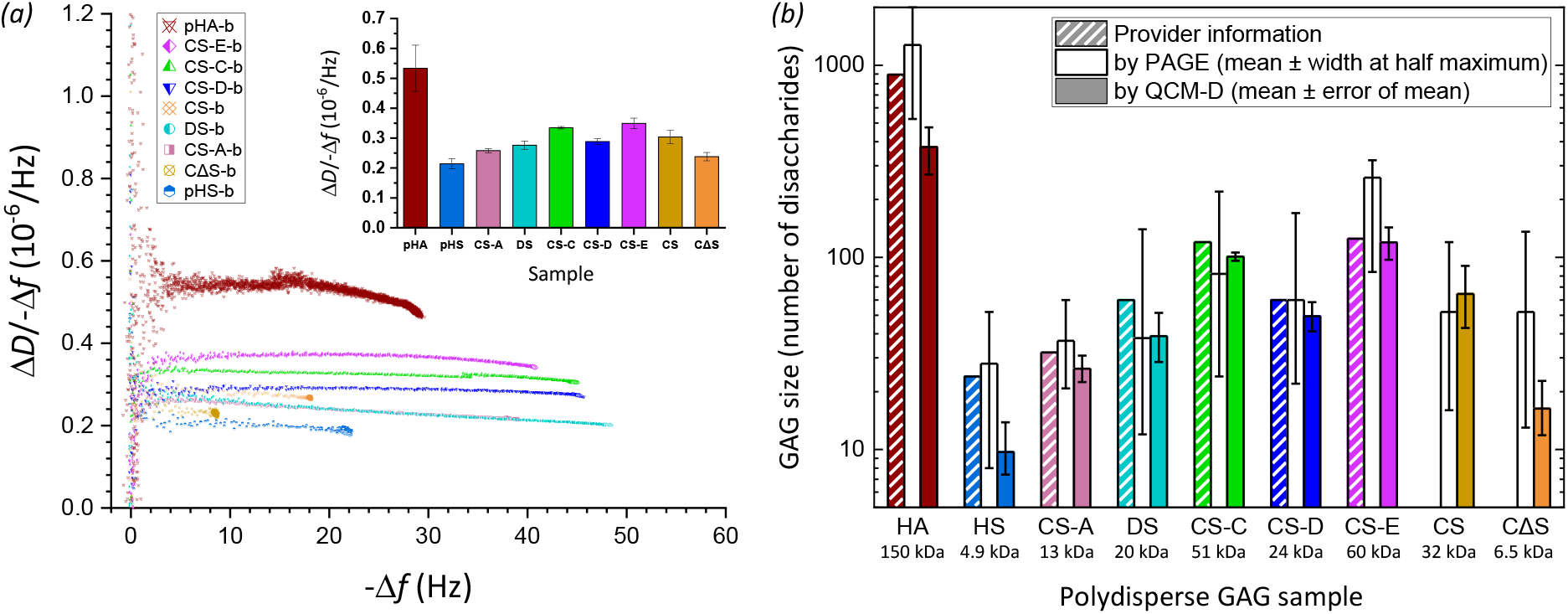
Application examples of chain sizing natural, isolated GAG preparations. *(a)* Plot of the Δ*D*/-Δ*f* ratio *vs*. -Δ*f* (*i* = 3) for various polydisperse GAG-b polysaccharide preparations of HA and other GAG types (chondroitin sulfate – CS, CS-A, CS-C, CS-D and CS-E, chemically desulfated CS – CΔS, dermatan sulfate – DS, heparan sulfate – HS; symbols and colour codes as indicated). Original QCM-D data are shown in **Figure S9**. The inset shows the Δ*D*/-Δ*f* values at -Δ*f* = 2.5 Hz, as mean ± standard deviations for -Δ*f* values ranging between 2 and 3 Hz. *(b)* Comparison of the effective size (in number of disaccharides *n*_ds_) of surface-grafted GAG-b polysaccharides (as determined by QCM-D; *filled bars*) with the effective size distribution of GAG-b polysaccharides in solution (as determined by PAGE; *unfilled bars*) and the size estimates of the original GAGs given by the provider (bars with diagonal lines; taking disaccharide masses to be 400 Da for HA, and 500 Da for all sulfated GAGs). To calculate GAG sizes from QCM-D data, the Δ*D*/-Δ*f* values at -Δ*f* = 2.5 Hz of the polydisperse GAG-b preparations were compared with the standard curve (**Figure 4** and **Equation 1**). To calculate the effective size distribution from PAGE data, the migration distances of the peak intensity and the half-maximal intensities above background were compared to the migration distances of size-defined HA standards (**Figure S8**). The effective molecular masses of surface-grafted GAGs (in kDa) are given below the GAG names, and were derived from *n*_ds_ with disaccharide masses of 400 Da for HA and CΔS, and 500 Da for all sulfated GAGs.

Unlike HA, sulfated GAGs isolated from mammalian system are usually less than 100 kDa in size. To test if the developed method could size other GAGs with shorter lengths, we also analysed several sulfated polydisperse GAG polysaccharides derived from animal tissues from commercial sources, including heparan sulfate (HS), dermatan sulfate (DS), and chondroitin sulfate (CS) preparations purified with varying degrees of sulfation (CS-A, CS-C, CS-D and CS-E) (**Table 2**). The QCM-D results (**Figure S9** and **Figure 6a**) show that the trends for the calculated mean molecular masses are in good agreement with company estimates (**Figure 6b**) and PAGE analyses (**Figure S8b-c**), and thus confirm that the established standard curve (**Figure 4**) can also be used to determine the effective mean size of sulfated polydisperse GAGs on surfaces. A general tendency towards lower mean GAG sizes on surfaces can be observed, but the differences for the sulfated GAGs are typically less pronounced than for the polydisperse HA (**Figure 6b**). Since the sulfated GAGs were <100 kDa, steric hindrance by the forming brush should be negligible, and the reduction in mean size of these GAGs on surfaces is most likely due to faster diffusive transport of smaller molecules. An unusually large decrease in mean size is noticeable for the HS sample, and we suggest that this might be due to the exceptionally large size distribution of this particular sample: in PAGE, this particular pHS sample spread completely to the bottom of the gel (**Figure S8b**).

Lastly, we demonstrate that the method can be uniquely used to analyse how chemical treatment affects GAG size. In this particular example, the chondroitin sulfate (CS) preparation was chemically treated to produce desulfated chondroitin sulfate (CΔS; see Supplementary Methods for details). Comparative QCM-D analysis of the sample before and after chemical modification (and after the required conjugation with biotin at the reducing end; **Figure S9b** and **Figure 6a**; orange diamond and yellow circle with crosses, respectively) revealed that the desulfation process significantly reduced the mean GAG size (**Figure 6b**), revealing undesired fragmentation in the process. Of note, this type of comparative analysis is challenging with gel electrophoresis because the change in charge upon desulfation substantially affects the migration behaviour in addition to the effect of GAG size. Indeed, the CΔS and CS samples were virtually indistinguishable when analysed by gel electrophoresis (**Figure S8c**), most likely due to a coincidental cancellation of charge and size effects on the migration rate.

## 4. Discussion

### 4.1 Workflow and benefits of GAG sizing on surfaces by QCM-D

Using a large spectrum of size-defined HA polymers, we have established a method to quantify the mean size of GAGs grafted with one end to a planar surface. The method relies on QCM-D as the sole analysis technique, and exploits the monotonic increase of the Δ*D*/-Δ*f* ratio with GAG size. The standard curve of Δ*D*/-Δ*f vs*. GAG size, established here with HA in 150 mM NaCl for the third overtone at a set frequency shift of -Δ*f* = 2.5 Hz (**Figure 4** and **Equation 1**), provides good size sensitivity up to 500 kDa and can be applied for HA as well as other GAG types (**Figures 4** and **6**). This technique is especially useful for its ability to measure the mean GAG size directly at surfaces, which is particularly important for large polydisperse GAGs where the surface-attachment process can substantially modulate the size distribution, causing smaller chains to preferentially bind compared to the original size distribution in solution (**Figure 6**).

To facilitate the adoption of the method, we have prepared a workflow which recapitulates the main steps involved in GAG sizing on surfaces using this method (**Figure 7**). It comprises: (i) conjugation of an anchor tag (here biotin) to one end (here the reducing end) of the GAG chains, (ii) monitoring of GAG grafting to a planar and quasi-rigid surface by QCM-D (at 150 mM NaCl), (iii) determining the time-averaged Δ*D*/-Δ*f* ratio at -Δ*f* = 2.5 Hz from the parametric Δ*D*/-Δ*f vs*. -Δ*f* plot (*i* = 3), and (iv) determination of the mean GAG size with the aid of the standard curve (**Figure 4** and/or with **Equation 1**). We note that only the GAG binding data up to Δ*f* = -3 Hz are required for the GAG sizing to work. The GAG incubation process therefore can be shortened compared to the example shown in **Figure 7** thus reducing time and sample where this is desirable. An extended binding range though has the benefit of permitting further quality control, as the trends in the parametric Δ*D*/-Δ*f vs*. -Δ*f* plot can be compared with size-defined standards.

**Figure 7.**
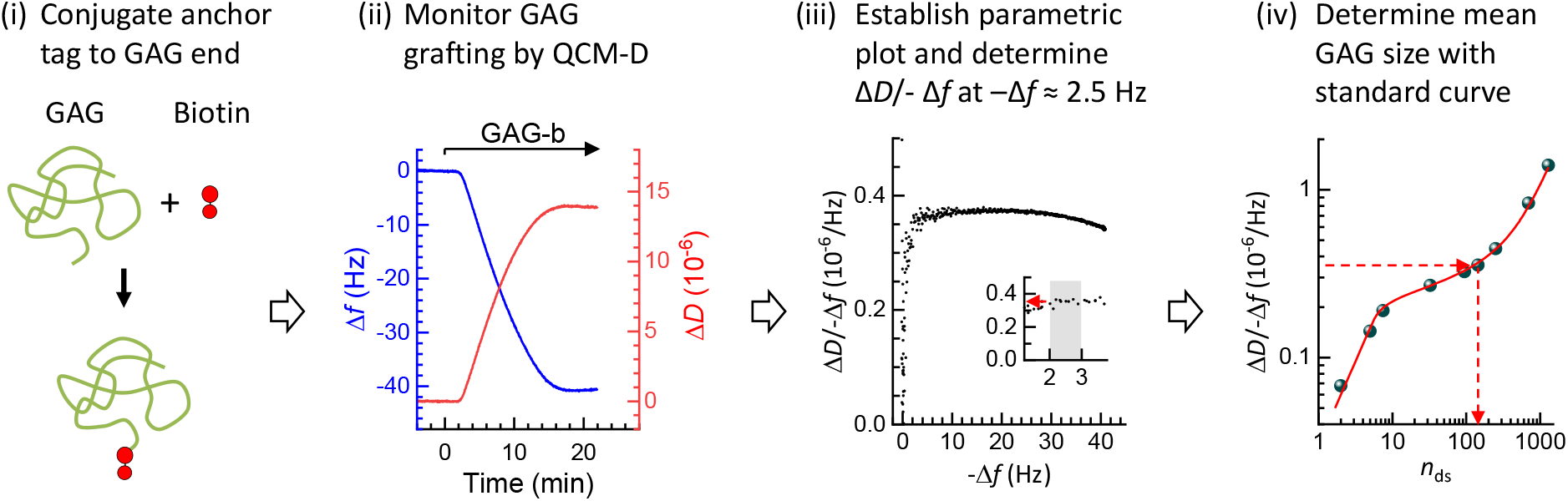
Flow diagram of the process of sizing surface-grafted GAGs. The four main steps are illustrated.

We expect that the here-established method will be most useful for the characterisation of GAG-based surface coatings. Benefits in this regard are that the GAG size is measured directly at the surface, and that the required data can be obtained non-destructively whilst monitoring GAG film formation by QCM-D. The latter is useful (*e*.*g*., for quality control) whenever GAG-functionalised surfaces are to be used further, *e*.*g*., for biomolecular interaction studies by QCM-D or other ‘solid-phase’ analysis techniques such as surface plasmon resonance or ellipsometry.

This QCM-D method should be of particular interest for the biophysical characterisation of GAGs isolated from tissues. Selective biotinylation of GAGs at their reducing end is well established [52]. The amounts of GAG required for size analysis are relatively small (a few μg or less) and would be achievable from tissue samples. Further on-surface analysis is possible to probe for selective interactions with GAG-binding proteins for glycobiological analyses. GAGs have also been widely exploited for biomedical applications such as implant coatings, biomaterial scaffolds and as nanoparticles for drug delivery. The insights and data from this study should be useful for improved characterisation of such materials, and for the design and manufacture of GAG-based coatings with improved performance for tissue engineering and medical device applications.

### 4.2 Sensitivity of the here-established method of GAG analysis

In Section 3.4, we established the slope α of the standard curve as a simple way to assess the size sensitivity of our GAG sizing method. Similarly, from the logarithmic presentation in **Figure 5**, it can be concluded that the HA size sensitivity improves at low ionic strength for oligosaccharides and smaller polysaccharides up to 58 kDa (*n*_ds_ ≈ 150). Although this benefit comes at the expense of a complete loss in size sensitivity for HA >58 kDa, this example illustrates that tuning of ionic strength could potentially be exploited to optimize GAG sizing sensitivity for certain applications.

Moreover, with the QCM-D response being sensitive to GAG size and GAG charge (**Figure 5**), it is conceivable that a screen of Δ*D*/-Δ*f* over a range of ionic strengths could in the future be employed to analyse the (mean) charge in addition to the (mean) size of surface-grafted GAGs. Such a methodology would benefit from a wider range of size- and charge-defined GAGs to establish a suitable ‘two-dimensional’ standard than what is currently accessible, although the compensation of charge and ionic strength [64, 70] can potentially also be exploited.

### 4.3 The complex shape of the standard curve reveals three distinct GAG film conformations

The shape of the here-established standard curve (**Figure 4**) is remarkably complex. Considering again a double logarithmic plot of the HA data (**Figure 8a**), four size-sensitivity regimes can be discerned. Regimes I and II are described by distinct power laws (**Figure 8a**, dashed red lines with slopes indicated), with rather well-defined slopes (*i*.*e*., powers) of 1.0 and 0.2, respectively. Regime IV may be the start of a plateau (power = 0) or close to a maximum (as more clearly seen with higher overtones, see **Figure S5**). In regime III, the sensitivity of Δ*D*/-Δ*f* for HA size is not described by a power law but instead gradually increases, and clearly is higher than in the neighbouring regimes II and IV.

**Figure 8.**
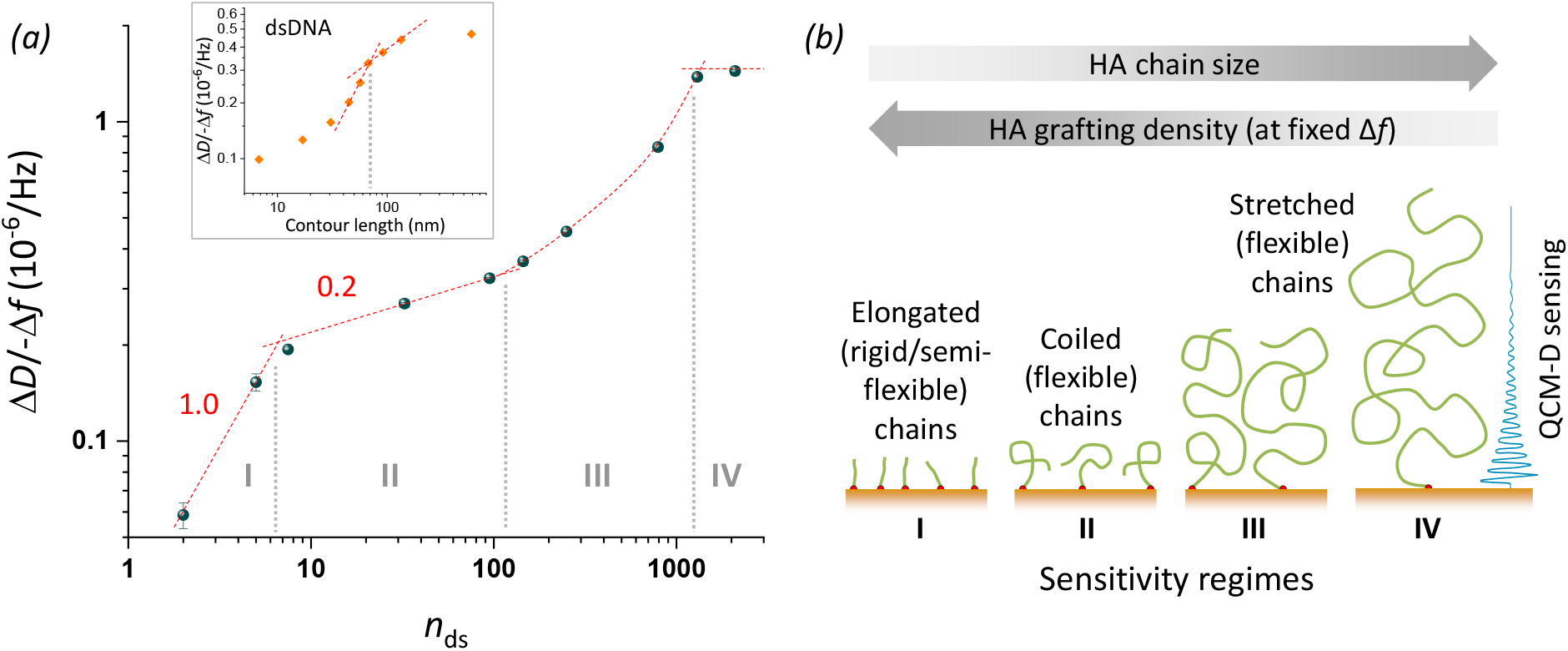
Sensitivity regimes of HA and underpinning mechanisms. *(a)* Double-logarithmic plot of the Δ*D*/-Δ*f* ratio (*i* = 3) at low HA surface density (-Δ*f* = 2.5 Hz) as a function of the mean number of disaccharides, *n*_ds_. Data for HA (blue spheres with error bars) are reproduced from **Figure 4**. Sensitivity regimes I to IV are separated by vertical grey dotted lines. Regimes I and II are discerned by their distinct power-law dependencies (red dashed lines with slope illustrate the approximate power); regime III does not obey a power law (red curved dashed line illustrates the trend) and regime IV is close to a plateau/maximum (indicated by a horizontal line). The inset shows equivalent data for dsDNA extracted from Figure 3 of Ref. [46], with two selected power law dependencies (dashed red lines). *(b)* Illustration of the proposed model to explain the observed sensitivity regimes. Both HA chain size and grafting density (which decreases with increasing chain size at the set -Δ*f* = 2.5 Hz) influence chain conformation, and distinct chain conformations are associated with sensitivity regimes I, II and III; ultimately, chain stretching exceeds the QCM-D sensing depth leading to a Δ*D*/-Δ*f* maximum and vanishing size sensitivity in regime IV.

What are the physical mechanisms giving rise to four distinct regimes? A comprehensive theoretical model that links GAG size to Δ*D*/-Δ*f* ratio remains elusive. However, the location of the boundaries between regimes hints at the regimes reflecting differences in HA chain conformation, as illustrated in **Figure 8b**. Specifically, the transition between regimes I and II is located at 7 disaccharides, equivalent to a contour length of *L*_c_ ≈ 7 nm. This value is comparable to the persistence length of HA (*L*_p, HA_ ≈ 4 nm), implying that rigid (*L*_c_ < *L*_p_) and semi-flexible (*L*_c_ ≈ *L*_p_) HA chains are probed in regime I, whereas flexible (*L*_c_ > *L*_p_) HA chains are probed in all other regimes. Tsortos et *al*. [46] have previously presented an analysis of Δ*D*/-Δ*f* values for one-end grafted double-stranded DNA (dsDNA), which is reproduced in **Figure 8a** (inset) for comparison. Although other aspects of the curve shape for dsNA are different, we note a similar transition from a stronger size dependence to a weaker size dependence at approximately 80 nm, again slightly larger than the persistence length of dsDNA (*L*_p, dsDNA_ ≈ 50 nm). Tsortos et *al*. [46] further suggested a theoretical model for the regime of rigid and semi-flexible chains that links the polymer shape (parametrised as intrinsic viscosity) to the Δ*D*/-Δ*f* values. With appropriate re-scaling based on this model, to account for differences in the mass per unit length and the diameter of the two polymers, we find that the data for HA and dsDNA superimpose rather well in the rigid chain regime (**Figure S10**). Taken together, these findings suggest that the transition between regime I (high size-sensitivity) and regime II (lower size sensitivity) is a general feature of polymer brushes and occurs when the contour length roughly matches the persistence length.

The origin of the transition between regimes II and III is less obvious. We propose that it relates to the onset of repulsion between grafted chains. As the chain size increases, the radius of gyration (which defines the mean size of the random coil formed by flexible HA chains in solution; *R*_g_) also increases. HA chains will retain their characteristic random coil conformation upon surface grafting as long as the radius of gyration is inferior to the mean anchor spacing (*d*_rms_ ≥ *R*_g_), but inter-chain repulsion at *d*_rms_ < *R*_g_ leads to chain stretching and film thickness changes. The first data point clearly departing from power 0.2 behaviour corresponds to an HA size of 250 disaccharides (*L*_c_ ≈ 250 nm; *M*_W_ ≈ 100 kDa). The radius of gyration for an HA chain of 250 nm contour length is *R*_g_ ≈ 21 nm [71]. We do not know what the exact anchor spacing of such an HA chain would be at Δ*f* = -2.5 Hz, but in light of attainable anchor spacings for very short HA chains (*d*_rms_ ≈ 5 nm for oligosaccharides) and very long HA chains (*d*_rms_ > 50 nm for HA 838 kDa; see section 3.2), a value in the range of 20 nm (*i*.*e*., consistent with a transition from random coil to brush conformation at this HA size) appears entirely reasonable.

Last but not least, the transition between regimes III and IV is likely defined by the limited sensing depth of QCM-D. The sensing depth in water (as well as ultrasoft HA films) is on the order of a few 100 nm [38, 72] which is rather close to the radius of gyration of the largest HA chains (*R*_g_ ≈ 75 nm for HA 838 kDa); therefore, even with only moderate chain stretching the HA brush thickness can readily exceed the QCM-D sensing depth.

Taken together, the different HA conformation regimes combined with the limited QCM-D sensing depth (**Figure 8b**) provide a first plausible explanation for the complex shape of the standard curve and the four distinct sensitivity regimes. In the future, further analysis with complementary techniques to directly measure HA grafting densities and film thicknesses may help to confirm and refine this model, and the insight gained could be exploited to further maximise size sensitivity for selected size ranges.

## 5. Conclusions

We have established an on-surface technique, based on QCM-D, to quantify the mean size of one-end grafted GAGs. The standard curve of Δ*D*/-Δ*f vs*. GAG size, shown in Figure 4 and represented by **Equation 1**, enables determination of the effective mean GAG size up to 500 kDa. The method is accurate for unsulfated GAGs such as HA, and also provides robust size estimates for sulfated GAGs such as HS and CS, with a typical resolution below 10%. By systematically analysing GAG brushes as a function of GAG size and type, this work has also provided new insight into basic physical properties of such brushes, such as the kinetics of GAG brush formation, the stability of GAG brushes, and how GAG conformation in GAG brushes is sensed by QCM-D. We have illustrated the importance of measuring the mean GAG size directly at surfaces, in particular for large polydisperse GAGs, as the surface-attachment process can substantially reduce the mean size compared to the mean size of the GAG pool in solution. The GAG sizing method will be most useful for the characterisation of GAG-based surface coatings, and should be of particular interest for the biophysical characterisation of GAGs isolated from tissues, and for the design and quality control of GAG-based coatings with improved performance for tissue engineering and medical device applications.

## Supporting information

Supplementary methods, figures and references

## Acknowledgements

We thank Anne Geert Volbeda and Jeroen Codee for kindly providing HAdp15-b, Hugues Lortat-Jacob for kindly providing HS polysaccharides and oligosaccharides for biotinylation, and Dhruv Thakar for biotinylating the polydisperse HA and HS polysaccharides, and the HS oligosaccharides.

This work was supported by an Oklahoma Center for Advancement of Science and Technology grant (HR18-104, to PLD), a Long Duration Visitor Grant from Labex Tec21 (ANR-11-LABX-0040, to DD and RPR), a Research Grant from the BBSRC, UK (BB/T001631/1, to RPR), the Leverhulme Trust (RPG-2018-100, to JCFK and RPR), an EPSRC CDT project funding (EP/L014823/1, to LS and JCFK), and the Center of Reconstruction Neuroscience -NEURORECON (CZ.02.1.01/0.0/0.0/15_003/0000419 to JCFK).

## Author contributions

SS designed experiments, collected and analysed data, and drafted the original manuscript. LS conceived the study, designed experiments, and collected and analysed data. AG, DEG, JD and PLD provided essential reagents. LD provided essential reagents and collected data. ARER and XZ collected data. DD analysed data. JCFK and RPR conceived the study, designed experiments, analysed data, drafted the original manuscript, supervised the project and raised funds. All authors contributed to manuscript review and editing.

