## Supplementary methods, figures and references for "A quartz crystal microbalance method to quantify the size of hyaluronan and other glycosaminoglycans on surfaces"

**Table of contents**

Supplementary methods S2

Supplementary figures S6

Supplementary references S19

**Supplementary methods**

***Biotinylation of GAGs***

All polydisperse GAGs were biotinylated (at their reducing end) by oxime ligation. pHA and HS were biotinylated as described previously [1]. In these two cases, a custom-made biotin derivative with an oligo(ethylene-glycol) ($EG_3$) linker and an oxyamine reactive group was used, thus adding an =N-O-$CH_2$-CO-NH-$(CH_2)_2$-$EG_3$-CH-NH-CO-$(CH_2)_4$-biotin moiety to the $C_1$ of the GAG's reducing-end GlcNAc.

CS-A, CS-C, CS-D, CS-E, CS, CΔS and DS polysaccharides (see Table 2 for details of these samples) were biotinylated following the protocol described in ref. [1], with modifications. A slightly different biotin derivative (EZ-link alkoxyamine PEG4-biotin; Thermofisher Scientific) was used here, resulting in a =N-$EG_4$-NH-CO-$(CH_2)_4$-biotin moiety at the $C_1$ of the GAG's reducing-end. Reaction conditions were: 5 mg·ml$^{-1}$ GAG, 10 mM EZ-link Alkoxyamine PEG4-biotin, 20 mM aniline (Sigma Aldrich), 50 mM acetate buffer, pH 4.5, 37 °C, vortex at 300 rpm overnight. The target polysaccharide was subjected to dialysis against u$H_2O$ through a cellulose membrane with a 3.5 kDa molecular weight cut-off (Thermofisher Scientific). All biotinylated GAGs were stored at -20°C until use.

***PAGE densitometry***

Images of polyacrylamide gels with GAGs stained by Alcian blue were analysed to verify the quality of size-defined GAGs, and to estimate the mean and spread in size of polydisperse GAGs. Image J software was used to extract density profiles of all relevant lanes (including ladders with molecular mass markers) from the images. Two horizontal lines were drawn across the top and bottom of the resolving gel which defined the zero and maximum migration distance, respectively. The effective migration distance of GAG bands was characterised by the location of the peak in intensity, and the width of the band was operationally defined from the migration distances at which half-maximal intensity above background was observed. For size-defined GAGs (incl. ladders and standards), an algorithm to fit one or multiple Gaussian peaks (implemented in Origin software) was employed to extract the peak position and the width of the GAG bands. For polydisperse samples, a Gaussian did not reproduce the experimental data well and peak position and width were instead estimated manually.

***Quality control of surface functionalisation***

We used QCM-D to monitor the surface functionalisation prior to the formation of GAG brushes, see Figure 1c (0 to 50 min) for representative data. The shifts observed upon incubation of the silica-coated sensor surface with biotin-displaying small unilamellar vesicles (b-SUVs; Figure 1c, 7 to 17 min) were $\Delta f$ = -25 ± 1 Hz and $\Delta D$ < 0.3 × $10^{-6}$. These values are characteristic of formation of an SLB of good quality [2]. The transient extrema in frequency and dissipation shifts during the SLB formation process (Figure 1c, at around 11 min) are well known, and reflect the process of SUVs initially adsorbing intact and then rupturing and spreading to coalesce into a confluent lipid bilayer [2]. The additional shifts of $\Delta f$ = -26 ± 1 Hz and $\Delta D$ = 0.5 × $10^{-6}$ upon exposure of SLBs to SAv (Figure 1c, 27 to 33 min) indicate the formation of a thin and densely packed SAv monolayer [3].

***Quantitative analysis of QCM-D data - accounting for solution effects and experimental errors***

We systematically examined the influence of ionic strength on brushes of varying HA chain lengths, by intermittent exposures to 0 mM and 1000 mM NaCl, in addition to the standard 150 mM NaCl in which the brushes were formed (Figure 1c and Figure S2). For quantitative analysis, the QCM-D responses due to salt-induced changes in the HA brushes had to be separated from the QCM-D responses due to salt-induced changes in the viscosity and/or density of the buffer solution. This compensation was accomplished with the aid of control experiments, as follows.

*Accounting for solution effects on the QCM-D response.* The experiments provide frequency shifts $\Delta f^{\mathrm{con}}_{[\mathrm{NaCl}]}$ and $\Delta f^{\mathrm{HA}}_{[\mathrm{NaCl}]}$, and dissipation shifts $\Delta D^{\mathrm{con}}_{[\mathrm{NaCl}]}$ and $\Delta D^{\mathrm{HA}}_{[\mathrm{NaCl}]}$, at a suitable time *t* after HA brush formation. Superscript 'con' refers to control experiments with a streptavidin monolayer only (no HA brush), and superscript 'HA' refers to experiments with an HA brush. The subscript '[NaCl]' refers to the NaCl concentration (in units of mM) and is either 0, 150 or 1000. All shifts were measured relative to the QCM-D responses prior to HA brush formation (*i.e.*, for the streptavidin monolayer in 150 mM NaCl) which thus defined the baseline in each experiment. To minimise the impact of noise on the analysis, all frequency and dissipation shifts were averaged over a suitable time window (of typically 3 min duration) where the responses were stable. The standard error of the mean (SEM) of the temporal averages ($\Delta f < 0.05$ Hz and $\Delta D < 0.01 \times 10^{-6}$) was much smaller than the estimated errors due to baseline drifts, and was therefore neglected in the error analysis (*vide infra*).

Frequency and dissipation shifts due to the HA brush are:

$$\Delta f^{\mathrm{HA}}|_0 = \Delta f_0^{\mathrm{HA}} - \Delta f_0^{\mathrm{con}}, \qquad \Delta D^{\mathrm{HA}}|_0 = \Delta D_0^{\mathrm{HA}} - \Delta D_0^{\mathrm{con}} \qquad \text{(S1a)}$$

$$\Delta f^{\mathrm{HA}}|_{1000} = \Delta f_{1000}^{\mathrm{HA}} - \Delta f_{1000}^{\mathrm{con}}, \qquad \Delta D^{\mathrm{HA}}|_{1000} = \Delta D_{1000}^{\mathrm{HA}} - \Delta D_{1000}^{\mathrm{con}} \qquad \text{(S1b)}$$

The vertical bar on the left-hand side of the equations indicates that shifts are corrected for any solution effects. We further consider the following offsets:

$$\Delta\Delta f_0^{\mathrm{con}} = \Delta f_{150}^{\mathrm{con}} - \Delta f_0^{\mathrm{con}}, \qquad \Delta\Delta D_0^{\mathrm{con}} = \Delta D_{150}^{\mathrm{con}} - \Delta D_0^{\mathrm{con}}. \qquad \text{(S2a)}$$

$$\Delta\Delta f_{1000}^{\mathrm{con}} = \Delta f_{150}^{\mathrm{con}} - \Delta f_{1000}^{\mathrm{con}}, \qquad \Delta\Delta D_{1000}^{\mathrm{con}} = \Delta D_{150}^{\mathrm{con}} - \Delta D_{1000}^{\mathrm{con}}. \qquad \text{(S2b)}$$

These offsets can be readily extracted from experiments, as illustrated in Figure SM1 for the frequency shifts. From Equations (S1) and (S2), and setting $\Delta f_{150}^{\mathrm{con}} = \Delta D_{150}^{\mathrm{con}} = 0$ to correct for any potential baseline drifts in the control experiments, we obtain:

$$\Delta f^{\mathrm{HA}}|_0 = \Delta f_0^{\mathrm{HA}} + \Delta\Delta f_0^{\mathrm{con}}, \qquad \Delta D^{\mathrm{HA}}|_0 = \Delta D_0^{\mathrm{HA}} + \Delta\Delta D_0^{\mathrm{con}}. \qquad \text{(S3a)}$$

$$\Delta f^{\mathrm{HA}}|_{1000} = \Delta f_{1000}^{\mathrm{HA}} + \Delta\Delta f_{1000}^{\mathrm{con}}, \qquad \Delta D^{\mathrm{HA}}|_{1000} = \Delta D_{1000}^{\mathrm{HA}} + \Delta\Delta D_{1000}^{\mathrm{con}}. \qquad \text{(S3b)}$$

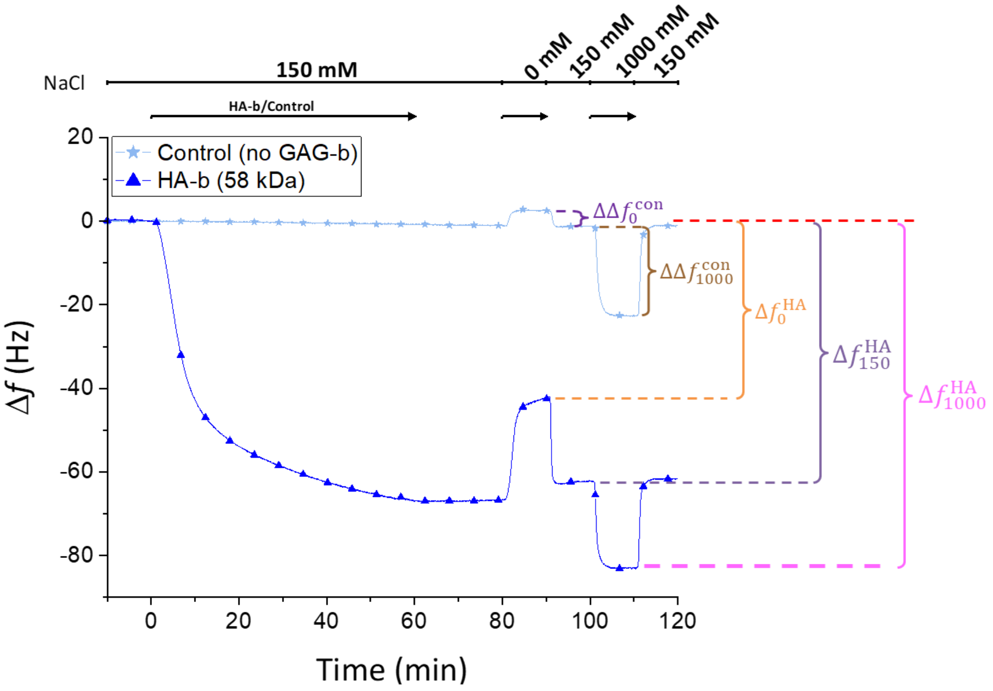


**Figure SM1. Illustration of salt-related offsets in QCM-D responses.** Frequency shift data are shown (taken from Figure S2) for a control experiment (*line with star symbols*), and for a brush formed with HA-b 58 kDa (*blue line with triangle symbols*). Offsets $\Delta\Delta f_0^{\mathrm{con}}$ and $\Delta\Delta f_{1000}^{\mathrm{con}}$, as well as shifts $\Delta f_0^{\mathrm{HA}}$, $\Delta f_{150}^{\mathrm{HA}}$ and $\Delta f_{1000}^{\mathrm{HA}}$ are indicated. Dissipation parameters were defined analogously. The dashed red line represents the 0 Hz baseline.

*Error analysis.* The typical time elapsing from the start of HA brush formation (where the baseline is determined) to the analysis of salt effects was approximately 90 min. Over such a long time, baseline drifts may occur at a level that far exceeds the experimental noise. From the control experiments, we estimated the magnitude of these drifts to lie within ±0.5 Hz in frequency, and within ±0.1 × $10^{-6}$ in dissipation, over 90 min. These uncertainties were associated with $\Delta f_{0/150/1000}^{\mathrm{HA}}$ and $\Delta D_{0/150/1000}^{\mathrm{HA}}$ and used for error propagation to reflect the limits of quartz sensor and equipment stability.

The offsets $\Delta\Delta f_{0/1000}^{\mathrm{con}}$ and $\Delta\Delta D_{0/1000}^{\mathrm{con}}$ were virtually unaffected by baseline drifts because the time elapsed between the measurement of shifts at 150 mM and either 0 mM NaCl or 1000 mM NaCl was short (< 10 min). We further improved on accuracy by averaging the offsets over several independent control experiments (*n* = 3) and included the SEM (0.08 Hz in frequency, and 0.04 × $10^{-6}$ in dissipation at 0 mM NaCl, and 0.15 Hz in frequency, and 0.07 × $10^{-6}$ in dissipation at 1000 mM NaCl, for *i* = 3) into the error propagation.

***Quantitative analysis of QCM-D data – determining errors in ΔD/-Δf at -Δf = 2.5 Hz***

Errors for Δ*D*/-Δ*f* at -Δ*f* = 2.5 Hz were determined as the time-averaged mean ± standard deviations of Δ*D*/-Δ*f* for -Δ*f* values ranging between 2 and 3 Hz. This simple approach worked well for all GAG sizes up to 800 disaccharides: in these cases, Δ*D*/-Δ*f* was virtually constant within the -Δ*f* = 2..3 Hz interval, and temporal drifts in Δ*D* and Δ*f* were negligible, implying that the temporal variations in Δ*D*/-Δ*f* are essentially due to data noise. For the two longest HA-b chains (520 KDa and 838 kDa), the Δ*D*/-Δ*f* ratio exhibited substantial changes across the -Δ*f* = 2..3 Hz interval (see Figure 3), and ~10 min or more of HA-b incubation were required to reach -Δ*f* = 2.5 Hz, implying systematic temporal variations and drifts could not be neglected. For these two HA sizes, we used a local linear interpolation of the -Δ*f vs.* time trace around -Δ*f* = 2.5 Hz to determine the time $t_{2.5\mathrm{Hz}}$ at which -Δ*f* = 2.5 Hz was reached, then used a local linear interpolation of the Δ*D vs.* time trace around $t_{2.5\mathrm{Hz}}$ to determine the corresponding dissipation shift, and finally, the Δ*D*/-Δ*f* ratio at -Δ*f* = 2.5 Hz. Temporal drifts (within rates of ±0.33 Hz/hour in frequency and ±0.067 × $10^{-6}$/hour in dissipation), were here included in the error estimation, in addition to the noise of the Δ*D vs.* time traces. This method worked well for overtones *i* = 3 and 5. For *i* = 7, the -Δ*f* values for HA-b 520 KDa and HA-b 838 kDa typically reached just ~2 Hz after 60 min of incubation. In these cases, the Δ*D*/-Δ*f vs.* -Δ*f* traces were extrapolated to 2.5 Hz by a local linear fit of all data -Δ*f* > 1.5 Hz.

***Desulfation of chondroitin sulfate***

CS (see Table 2 for sample details) was desulfated following a published procedure [4]. CS was stirred at 5.0 mg·mL$^{-1}$ in methanol (VWR) containing 10% *v/v* acetyl chloride (Merck). CS was centrifuged, and the acidic methanol solution was replaced on days one, three and seven of the seven day reaction, leading to formation of the methyl ester of chondroitin. The acidic methanol was aspirated off, and the product was dissolved in 20 mL of distilled deionised water per gram of starting CS, before being precipitated in an excess of ethanol (VWR). The precipitate was filtered, washed with cold ethanol and diethyl ether, and dried *in vacuo*. The dried product was demethylated at 25 mg·mL$^{-1}$ in 0.1 M potassium hydroxide (Thermofisher Scientific) for 24 h to give desulfated chondroitin. The product was neutralised in 4 mL of 100 mg·mL$^{-1}$ potassium acetate (Merck) in 10% *v/v* acetic acid (Thermofisher Scientific) per gram of starting product, and precipitated in an excess of ethanol. The CΔS product was filtered, washed with cold ethanol and diethyl ether, dried *in vacuo*, and stored at 4 °C until use. Desulfation was confirmed with the dimethylmethylene blue assay for sulfated GAGs [5] and infrared spectroscopy [6] (Figure SM2).

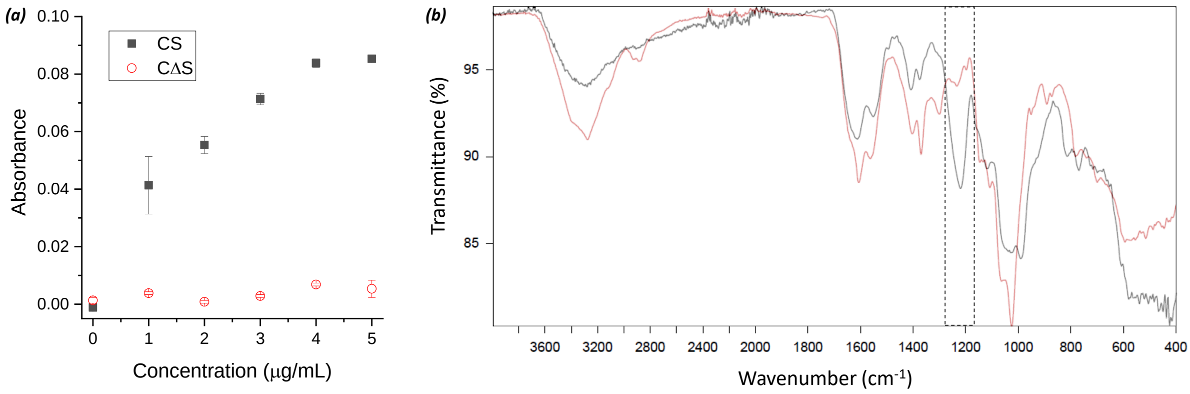


**Figure SM2. Analysis of desulfation of CS to CΔS**. (*a*) The dimethylmethylene blue (DMMB) assay confirms removal of the sulfate groups on CS. Standard curves ranging from 0 to 5 μg·mL$^{-1}$ of CS or CΔS were assayed by DMMB according to established protocols [5], and absorbance (above the background of the pure solvent) was measured at 525 nm (Envision plate reader, PerkinElmer); values shown are the mean ± S.D. for two technical replicates. Absorbance was virtually zero for the CΔS sample (*red*), and considering the absorbance for CS and experimental error we estimate that >90% of the sulfates were removed in CΔS. (*b*) Infrared (IR) spectroscopy of CS (black) and CΔS (red). Overlay of CS and CΔS show a characteristic sulfate peak (appearing as a trough in the transmission spectrum) at approximately 1200 cm$^{-1}$ [6] for CS, which is absent for CΔS (*dashed black box*).

**Supplementary Figures**

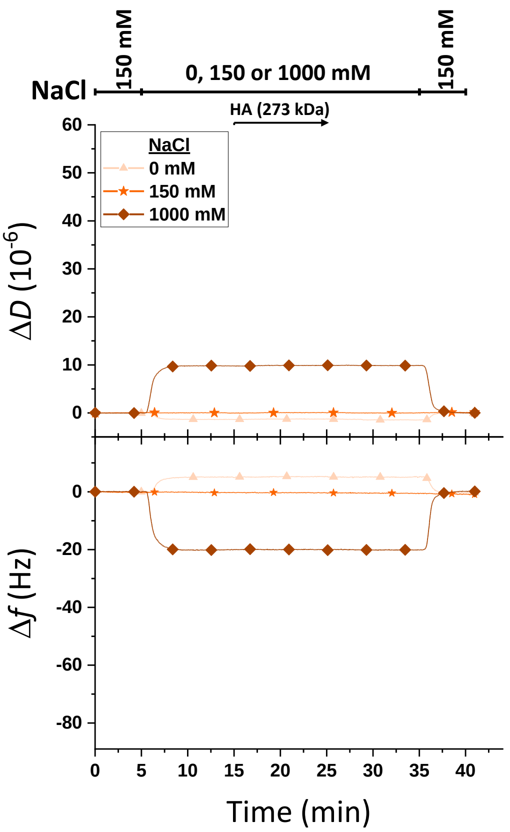


**Figure S1. HA-b is surface-anchored exclusively *via* the biotin at the reducing end.** QCM-D data are shown with incubation steps and NaCl concentrations indicated analogous to Figure 1c. A streptavidin monolayer was formed as shown in Figure 1c, and exposed to pH-buffered solutions with low (0 mM), close-to-physiological (150 mM) or high (1000 mM) NaCl concentrations followed by incubation of HA (273 kDa) that lacked a biotin tag. The lack of any measurable response upon HA incubation indicates that HA-b anchors to the streptavidin monolayer exclusively *via* the biotin tag at the reducing end, for all NaCl concentrations studied. Incubation conditions: buffer – 10 mM HEPES, pH 7.4, with NaCl as indicated; HA (273 kDa) – 20 µg/mL.

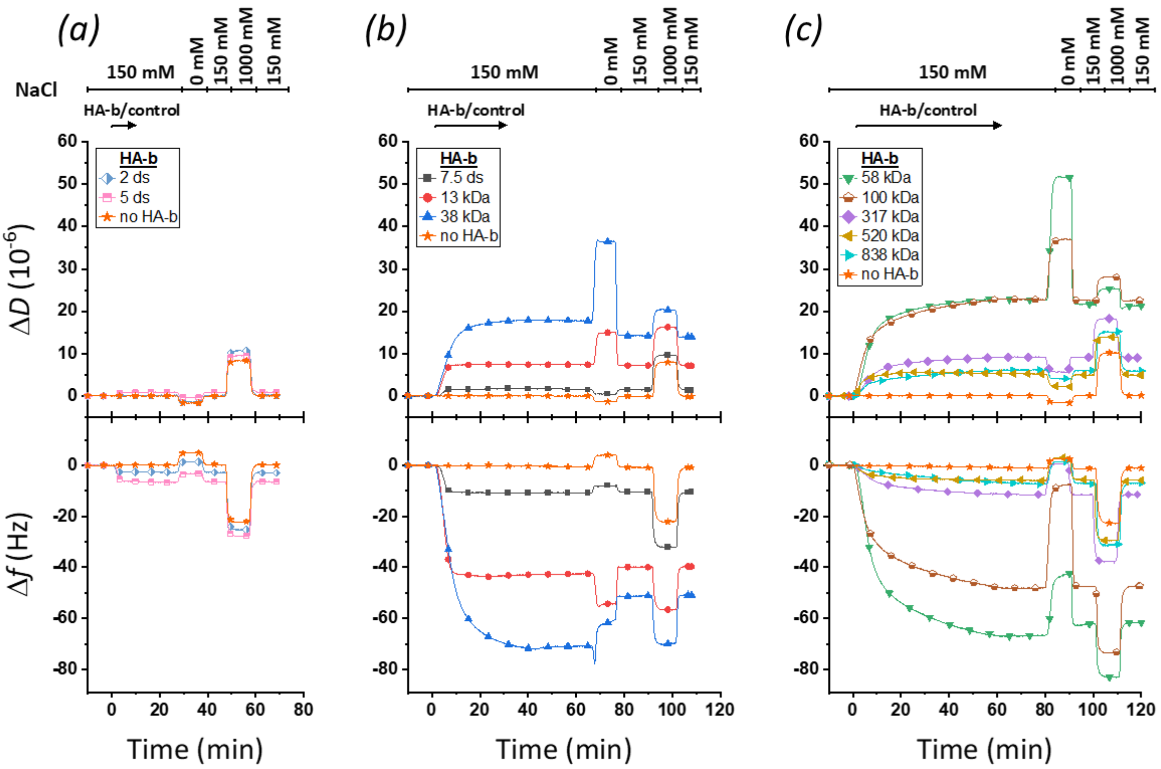


**Figure S2. QCM-D responses for brushes of size-defined HA.** Representative data (Δ*f* – *lower panels*; Δ*D* – *upper panels*; *i* = 3) for HA-b of defined sizes, ranging from oligosaccharides made from 2 disaccharides (ds) to polysaccharides of 838 kDa (*coloured lines with symbols*, as indicated), and for controls without HA-b (*orange lines with star symbols*). Experiments were performed on streptavidin monolayers, formed as shown in Figure 1c, and incubation steps and NaCl concentrations are indicated analogous to Figure 1c.

It can be seen that QCM-D responses for all HA polysaccharides are substantially affected by a reduction in ionic strength. A switch from 150 mM to 0 mM NaCl induced marked changes in Δ*f* and Δ*D*, although with varying trends depending on HA-b size: HA-b 13 kDa shows a net decrease in frequency and a net increase in dissipation, HA-b 317 kDa and larger HA-b chains show opposite effects, and HA-b 58 kDa and 100 kDa show net increases in both frequency and dissipation. We attribute all these effects to brush swelling owing to increased intra-chain and inter-chain electrostatic repulsion in the absence of salt [7, 8], although the swelling manifests in different ways in the QCM-D response due to the limited QCM-D sensing depth (which is a few 100 nm in aqueous solution [9]). The shortest HA polysaccharides (HA-b 13 kDa, $n_{ds} \approx 33$) have a contour length of 33 nm (1 nm per disaccharide; Figure 1a). In this case, the whole brush resides well-within the sensing depth (since the brush thickness cannot exceed the contour length) and the added solvent coupled into the brush upon swelling is effectively sensed as a decrease in frequency. For the longer HA chains, on the other hand, the brush thickness can exceed the sensing depth even at 150 mM NaCl, and swelling thus does not entail a further frequency decrease. We note in passing that films with a thickness comparable to the sensing depth can generate non-monotonous QCM-D responses owing to a phenomenon termed 'film resonance' [10]; whilst this effect shall not be considered in detail here, it is a potential contributor to the complexity of the trends observed. We note that a reduction of ionic strength had only little net effect on the QCM-D responses for HA oligosaccharides: as mentioned earlier, oligosaccharides are fairly stretched at physiological ionic strength, and any further brush swelling at lower ionic strength therefore is expected to be minimal.

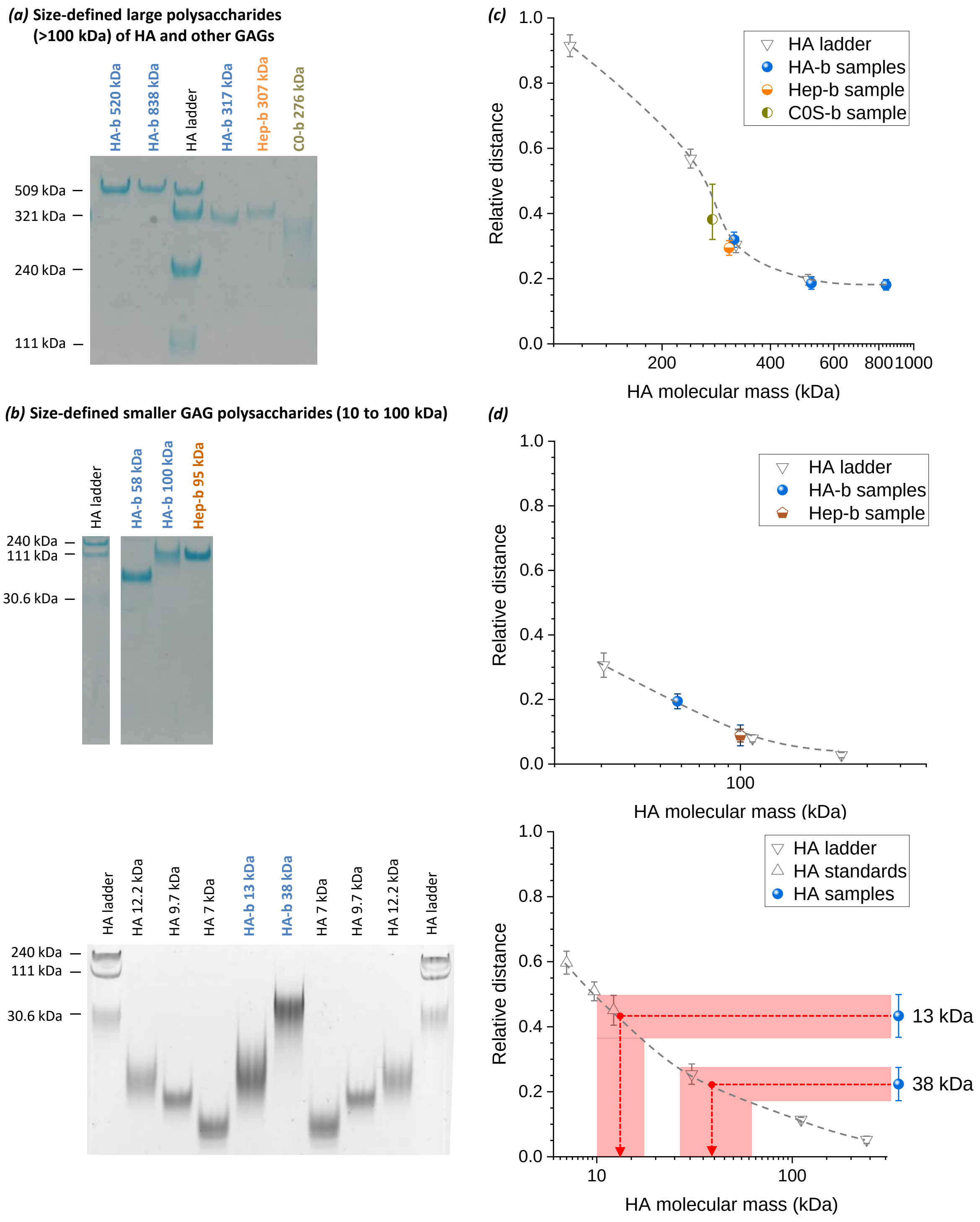


**Figure S3. Characterisation of size-defined GAG polysaccharides by polyacrylamide gel electrophoresis.** *(a)* Large polysaccharides (>100 kDa) of HA-b (*blue*), used to establish the standard curve (Figure 4, red line), and of additional GAGs (heparosan – Hep-b 307 kDa, *orange*; chondroitin – C0-b 276 kDa, *dark yellow*), used to confirm applicability of the standard curve (Figure 4). *(b)* Smaller polysaccharides (10 kDa to 100 kDa) of HA-b (*blue*), and of Hep-b 100 kDa (*orange*), which were also used to establish and confirm applicability of the standard curve, respectively. The indicated molecular masses of the polysaccharides were determined by size-exclusion chromatography multi-angle light scattering, except for the smallest HA-b sizes (13 and 38 kDa) which were estimated from comparison with size standards in the gel (see *bottom graph* in (*d*)). Ladders of size-defined HA, along with additional HA standards of 12.2 kDa, 9.7 kDa and 7 kDa (all non-biotinylated), are included for reference with molecular masses indicated. Conditions: gels contained 7% (*a*), 14% (*b*, top) or 8% (*b*, bottom) acrylamide; GAGs were stained with Alcian blue.

*(c-d)* Plots of the relative migration distance *vs.* the mean molecular mass, obtained by densitometry of the gels shown in (*a-b*). Mean values indicate the peak in staining intensity, and error bars the positions of half-maximal staining above background. Most data points fall on master curves (*grey dashed lines* are added as a guide to the eye; note that the data points for Hep-b 100 kDa and HA-b 100 kDa overlap), indicating that the mean molecular masses are close to their nominal values, and show consistent trends in peak width, indicating consistently narrow size distributions. Notable exception are C0-b 276 kDa, and to a minor extent HA-b 100 kDa, which show broader size distributions.

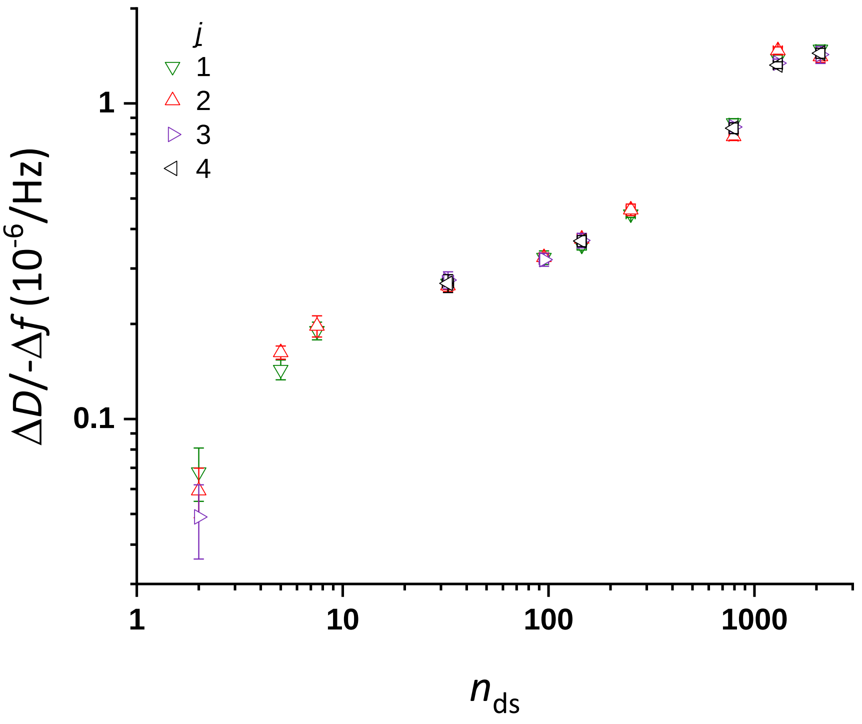


**Figure S4. Reproducibility of Δ*D*/-Δ*f* ratios.** Double logarithmic plot of Δ*D*/-Δ*f* (*i* = 3) at low GAG surface density (-Δ*f* = 2.5 Hz) as a function of the mean number of disaccharides ($n_{ds}$; mean ± 5% (errors are comparable to symbol widths), except for oligosaccharides which were taken to be pure in size, see **Table 1**), per HA-b chain. Data represent the mean ± error derived from individual experiments (see Supplementary Methods – Error analysis for details). Experimental repeats (*j* = 1 to 4, as indicated in the legend; 4 repeats for HA-b 13, 58, 317, 520 and 838 kDa, 3 repeats for HA-b 2 ds and 38 kDa, 2 repeats for HA-b 5 and 7.5 ds, and 100 kDa) for each HA-b size agree within their errors confirming good reproducibility.

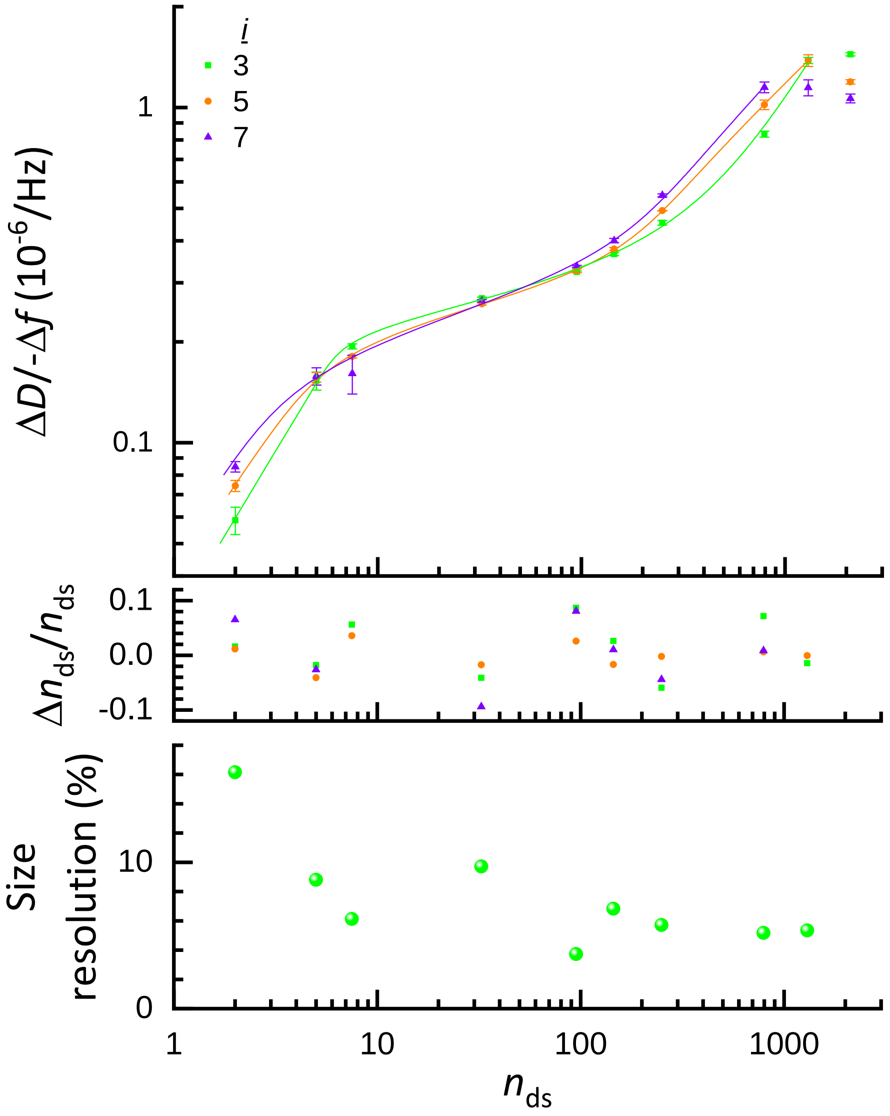


**Figure S5. The sensitivity of the $\Delta D/\text{-}\Delta f$ ratio to HA size depends only weakly on overtone.** (*Top*) Double logarithmic plot of $\Delta D/\text{-}\Delta f$ at low GAG surface density ($-\Delta f$ = 2.5 Hz) as a function of the mean number of disaccharides ($n_{ds}$) per HA chain, at selected overtones ($i$ = 3, 5 and 7; as indicated with symbol and colour code). Data represent the mean ± standard error across 4 (for HA-b 13, 58, 317, 520 and 838 kDa), 3 (for HA-b 2 ds and 38 kDa) or 2 (for HA-b 5 and 7.5 ds, and 100 kDa) experiments. For enhanced clarity, the uncertainties on $n_{ds}$ owing to finite HA size distributions are shown in Figure S4 but omitted here. The lines in matching colours interconnecting data points represent empirical fits to the data up to $n_{ds}$ = 1300 (520 kDa) for $i$ = 3 and 5, and up to $n_{ds}$ = 793 (317 kDa) for $i$ = 7 (*i.e.*, the region close to the plateau/maximum at the largest HA sizes where there is no clear size discrimination was not included in the fit). The formula corresponding to the lines are given as Equation (1) for $i$ = 3, as

$$n_{\text{ds}} = \frac{\Delta D}{-\Delta f} / \frac{10^{-6}}{\text{Hz}} \times \left(37.15 \frac{\text{Hz}}{10^{-6}} \frac{\Delta D}{-\Delta f} + 23.83\right) \frac{12.53 + \left(3.259 \frac{\text{Hz}}{10^{-6}} \frac{\Delta D}{-\Delta f}\right)^{-6.775}}{1 + \left(3.259 \frac{\text{Hz}}{10^{-6}} \frac{\Delta D}{-\Delta f}\right)^{-6.775}} \text{ for } i = 5, \quad \text{(S4a)}$$

and as

$$n_{\text{ds}} = \frac{\Delta D}{-\Delta f} / \frac{10^{-6}}{\text{Hz}} \times \left(17.50 \frac{\text{Hz}}{10^{-6}} \frac{\Delta D}{-\Delta f} + 20.20\right) \frac{16.99 + \left(3.125 \frac{\text{Hz}}{10^{-6}} \frac{\Delta D}{-\Delta f}\right)^{-5.128}}{1 + \left(3.125 \frac{\text{Hz}}{10^{-6}} \frac{\Delta D}{-\Delta f}\right)^{-5.128}} \text{ for } i = 7. \quad \text{(S4b)}$$

The slopes of the lines interconnecting data points are a measure for the relative size sensitivity of the Δ*D*/-Δ*f* ratio. Slopes are virtually independent of overtone number for oligosaccharides and small polysaccharides (up to $M_w$ = 38 kDa; $n_{ds}$ ≈ 100), and only minor differences are observed for the larger polysaccharides, indicating that size selectivity is similar for all overtones. A notable exception is the regime between 317 kDa ($n_{ds}$ ≈ 800) and 520 kDa ($n_{ds}$ ≈ 1300), where *i* = 3 shows very good size sensitivity, whereas *i* = 7 has already reached a maximum characteristic for the largest HA sizes; the earlier appearance of the maximum correlates with a reduced sensing depth expected for higher overtones [10, 11]. (*Middle*) Differences between the fit and experimental $n_{ds}$ values, normalised by the experimental $n_{ds}$ values. The residuals fall within the range of ±10%, except for $n_{ds}$ = 7.5 at *i* = 7, where the experimental spread in Δ*D*/-Δ*f* is very large. (*Bottom*) Size resolution (at *i* = 3) determined as $(\sigma/\mu)^{1/\alpha}$ from the slopes $\alpha = \mathrm{d}\ln(\Delta D/{-\Delta f})/\mathrm{d}\ln n_{\mathrm{ds}}$ according to Equation 1, and the standard deviations σ and mean values μ of Δ*D*/-Δ*f* measured across 2 to 4 experiments per sample (as specified above).

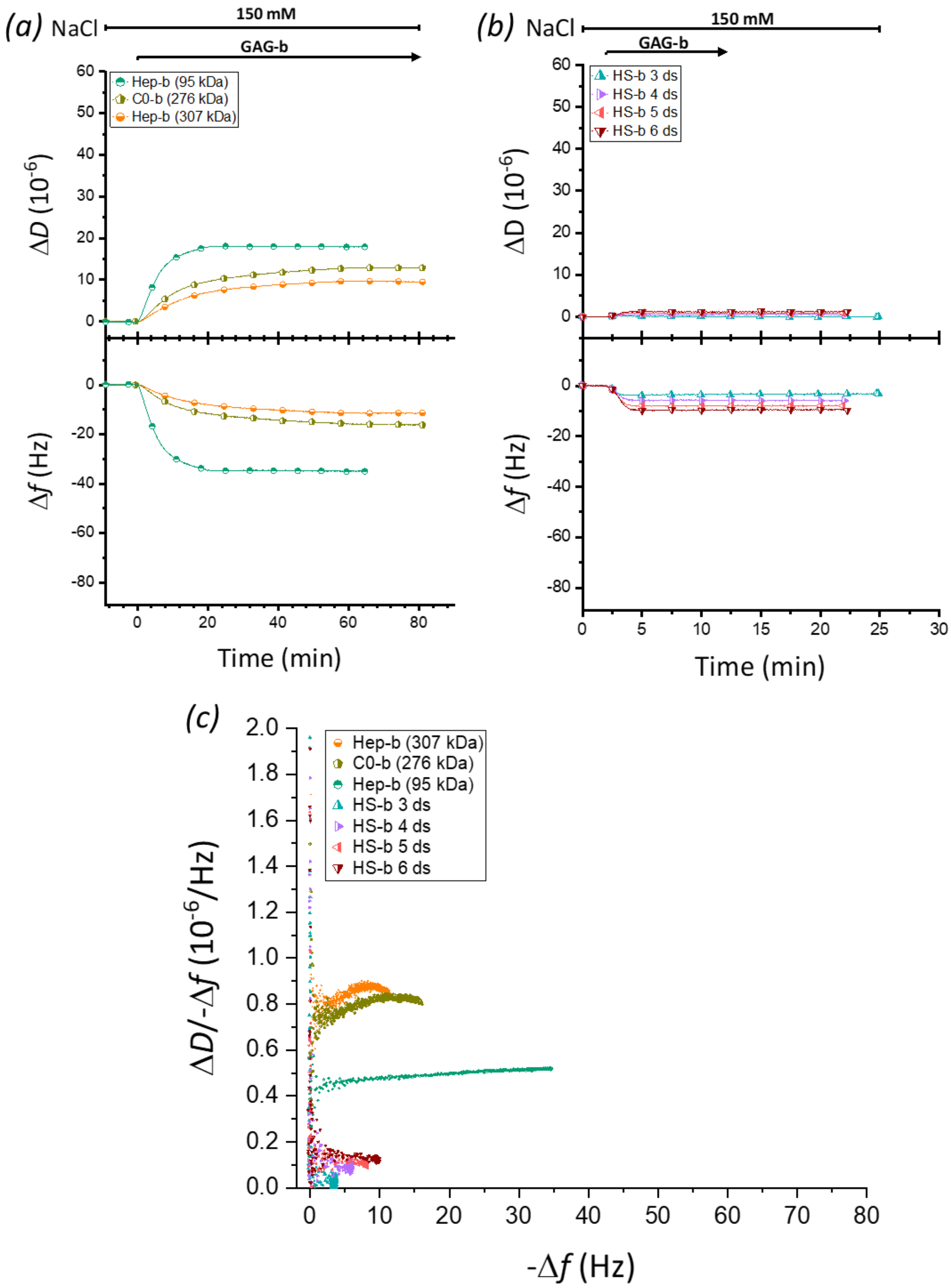


**Figure S6. QCM-D responses for brushes of other size-defined GAGs.** Representative data ($\Delta f$ – *lower panels*; $\Delta D$ – *upper panels*; $i$ = 3) for *(a)* Hep-b and C0-b polysaccharides and *(b)* HS-b oligosaccharides of defined sizes (as indicated). Data are displayed analogous to Figure 1c. Experiments with Hep-b and C0-b polysaccharides were performed on streptavidin monolayers, formed as shown in Figure 1c; data for HS-b oligosaccharides were taken from ref. [1]. *(c)* Parametric plot $\Delta D/\text{-}\Delta f$ *vs.* $-\Delta f$ ($i$ = 3) during the formation of brushes of size-defined Hep-b, C0-b and HS-b (as indicated with symbol and colour code). Incubation conditions: Hep-b and C0-b polysaccharides – 5 µg/mL; HS-b oligosaccharides – 50 µg/mL.

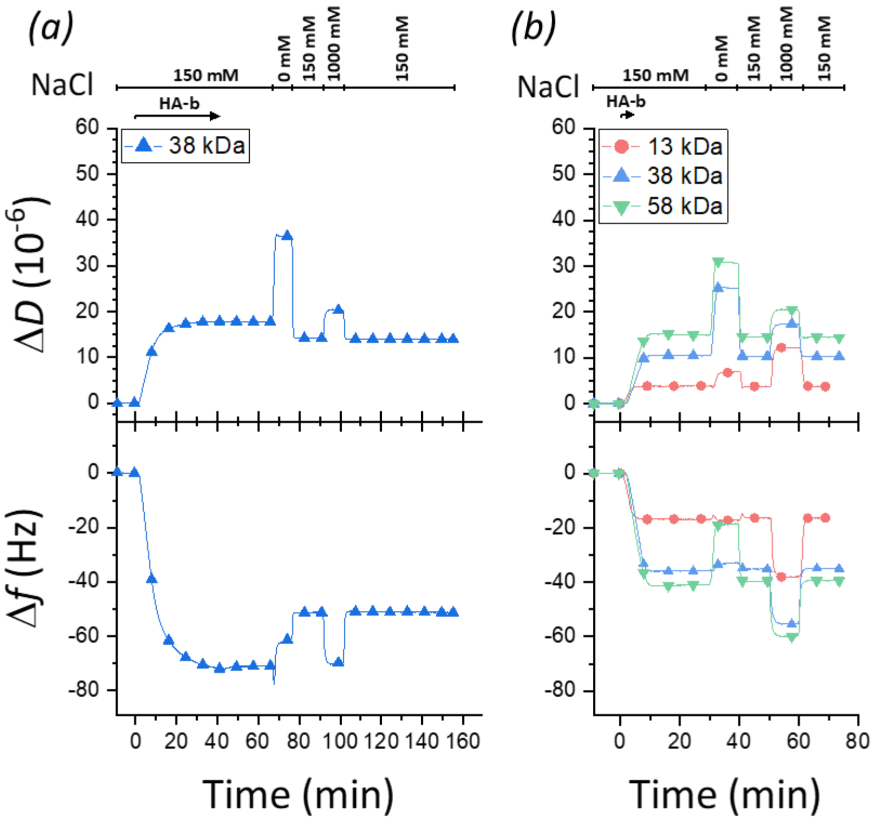


**Figure S7. Strong charge repulsion entails the release of HA from sufficiently dense brushes.** QCM-D data ($i$ = 3) are presented analogous to Figure 1c. *(a)* Responses for HA-b (38 kDa) in which HA brush formation was allowed to proceed to saturation ($\Delta f$ = -72 Hz) in 150 mM NaCl (reproduced from Figure S2b to facilitate comparison). It can be seen that $\Delta f$ increases during the 0 mM NaCl rinse (in contrast to the control which shows a decrease; see Figure 1c) and that the $\Delta D$ and $\Delta f$ responses at the end of brush formation are not fully recovered following a return to 150 mM NaCl after the 0 mM rinse (only $\Delta f$ = -51 Hz were retained after the low-salt quench). This indicates that a reduction in ionic strength triggers the release of HA from the surface. This effect was also observed for HA-b of 13 kDa (Figure S2b) and 58 kDa (Figure S2c). The enhanced repulsion between charged GAG chains at low ionic strength is expected to build up pressure within the brush which entails mechanical strain on the chain anchors. Our data suggest that, at the highest grafting densities, this strain is sufficiently high to trigger the release of HA chains from the brush. *(b)* Responses for HA-b (13 kDa, 38 kDa or 58 kDa) in which HA brush formation was interrupted by rinsing in buffer (150 mM NaCl) at an intermediate coverage ($\Delta f$ = -16 Hz, -36 Hz and -41 Hz respectively). Incubation conditions: HA-b (13 kDa) – 5 µg/mL, 3 min; HA-b (38 kDa) – 5 µg/mL, 5.5 min; HA-b (58 kDa) – 5 µg/mL, 7 min. The HA brushes were completely stable to a low salt exposure in these cases, indicating that chain release at low salt requires a minimal grafting density. It is notable that low salt did not drive any of the HA-b oligosaccharides out of the brush even though their grafting density was comparable to HA-b 38 kDa. Presumably, the small number of charges per chain in this case is insufficient to generate the required strain. On the other hand, the lack of chain release for HA-b polysaccharides (>58 kDa) is likely due to their much lower grafting density. Taken together, the data indicate that chain release requires low ionic strength and a combination of sufficiently high grafting density and chain size.

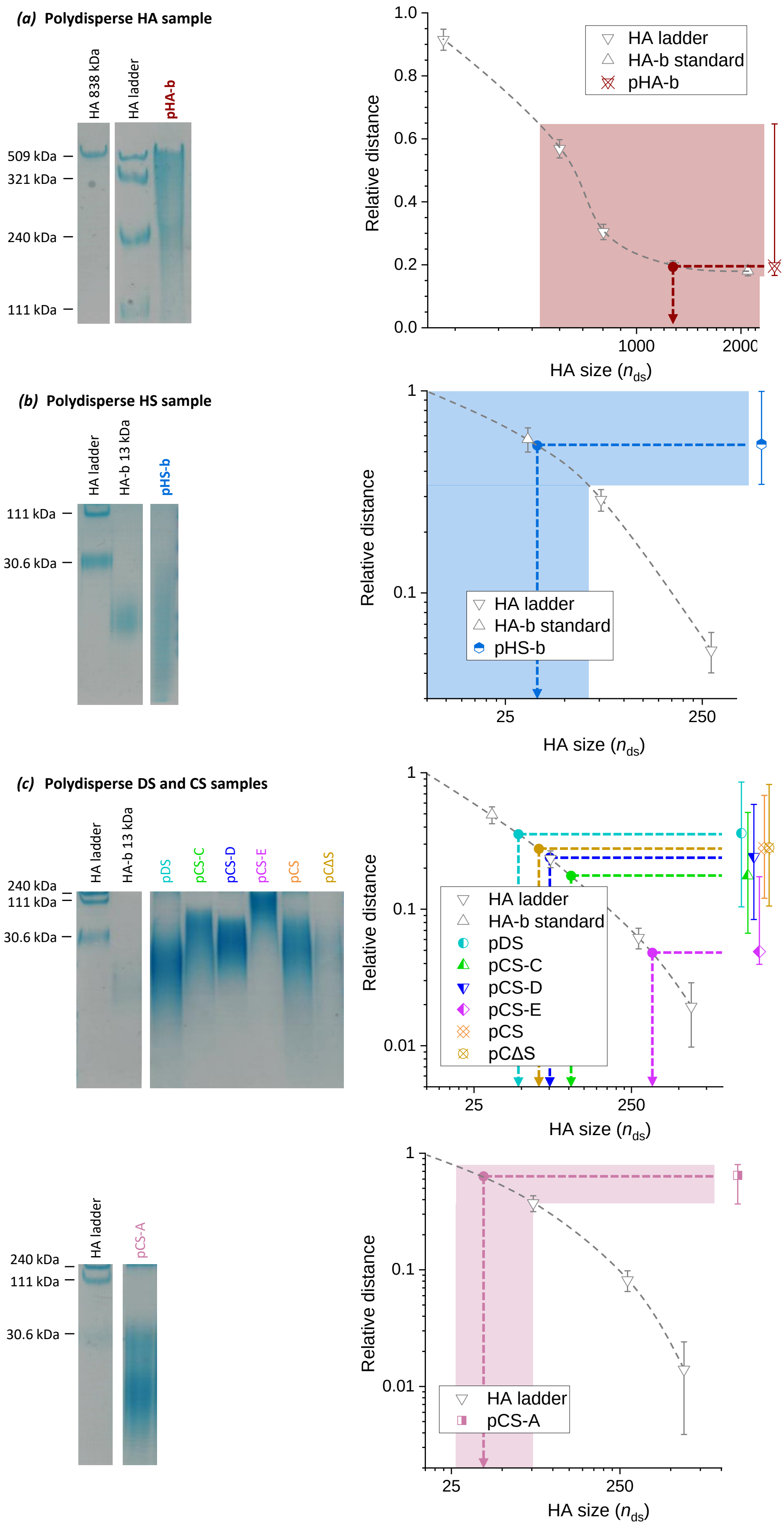


**Figure S8. Characterisation of polydisperse GAG polysaccharides by polyacrylamide gel electrophoresis.**

Reducing-end biotinylated GAGs of HA (*a*) and HS (*b*), and DS/CS prior to biotinylation (*c*) were used (see Table 2 for sample details). Data were analysed and are displayed as described in Figure S3, except for the shaded areas indicating errors of the size estimates which were omitted in one of the right graphs to enhance clarity. The HA size in the right graphs is expressed in number of disaccharides, $n_{ds} = M_W$ / 0.4 kDa. See Figure 6b for the results of the GAG size and size dispersity estimations based on HA standards. It should be noted that the migration distance is affected by the GAG charge density in addition to GAG size. This implies that the sizes of sulfated GAGs are underestimated because the sulfated GAGs have a higher charge density than HA and thus migrate faster at comparable size. Moreover, the degree of size underestimation will increase with the degree of sulfation, and sizing errors thus are expected to be largest for CS-D and CS-E. Conditions: gels contained 7% (*a*) or 14% (*b* and *c*) acrylamide; GAGs were stained with Alcian blue.

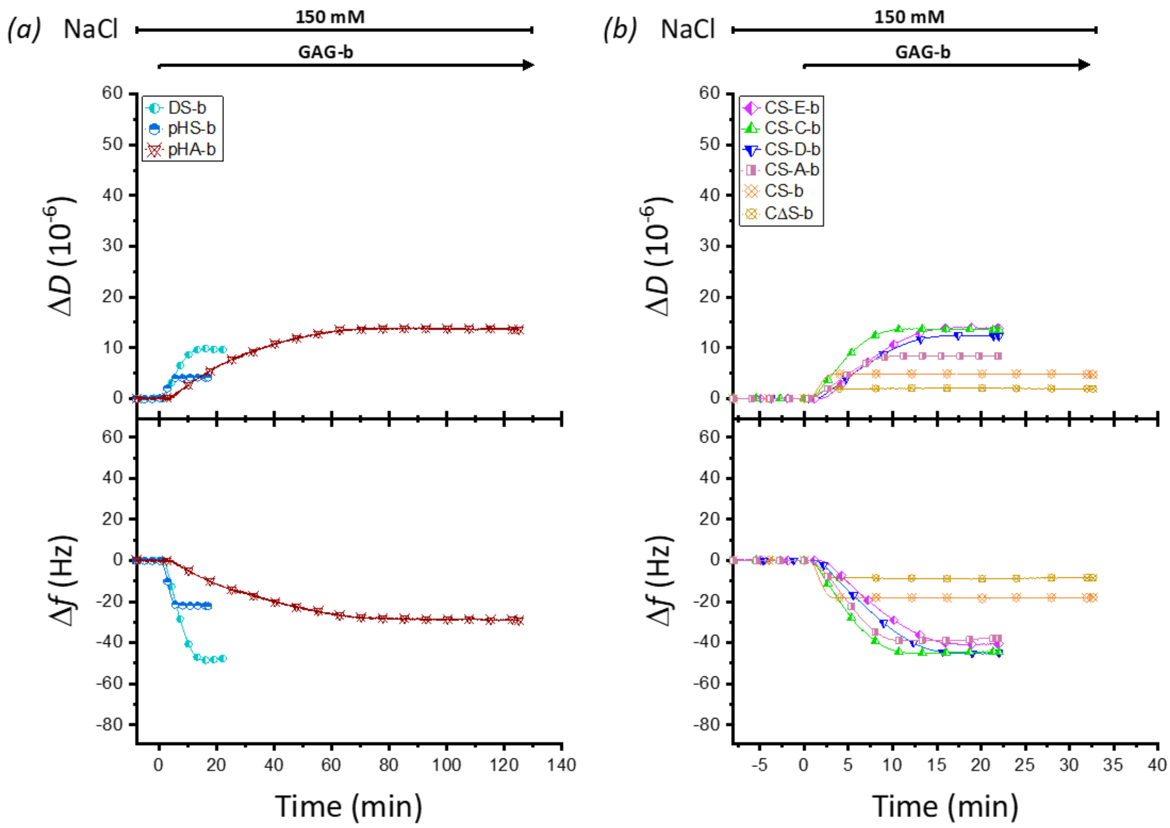


**Figure S9. QCM-D data for the formation of brushes with preparations of polydisperse GAGs.** (*a*) pHA, DS and pHS, and (*b*) CS GAGs (and the CΔS derivative) were biotinylated at their reducing ends, and incubated on streptavidin monolayers, formed as shown in Figure 1c. Representative data (Δ*f* – *lower panels*; Δ*D* – *upper panels*; *i* = 3) are shown analogous to Figure 1c. Incubation conditions: CS-A-b, CS-C-b, CS-D-b, CS-E-b, DS-b, CS-b, CΔS-b and pHS-b – 10 µg/mL; pHA-b – 50 µg/mL. Data for pHA-b was taken from ref. [1].

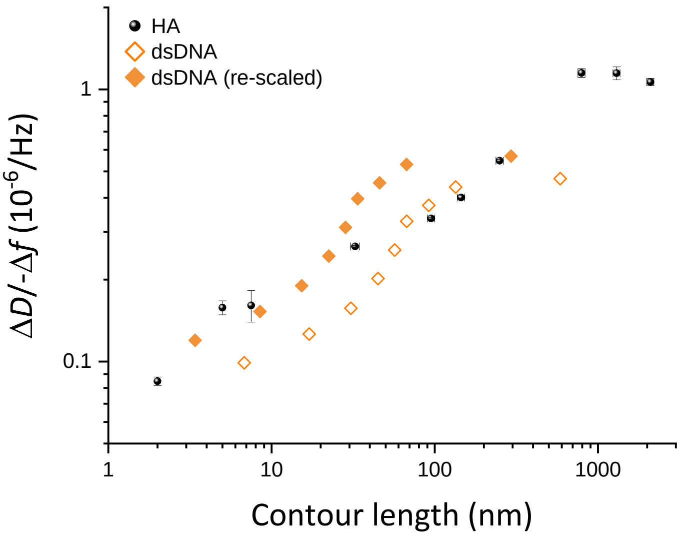


**Figure S10. Comparison of GAG and DNA brushes.** Data of Δ*D*/-Δ*f* for double stranded (ds) DNA (*i* = 7) were extracted from Figure 3 in ref. [12], and multiplied by the overtone number for frequency normalisation (*open orange diamonds*). In addition, these data were re-scaled (see below; *solid orange diamonds*) to account for the different mass per unit length and diameter of DNA *vs.* GAGs. The data for HA (*black spheres*) are presented analogous to Figure 4, but for *i* = 7 to enable direct comparison at this overtone, with the number of disaccharides converted to contour length (1 nm per disaccharide).

From Eq. 3 in ref. [12] and Eq. 3 in ref. [13], one can derive that $\Delta D/{-\Delta f} \approx [\eta] = v\frac{V_h N_A}{M_w}$, where $[\eta]$ is the specific viscosity of the solute. For a rigid rod (modelled as a prolate ellipsoid), the shape factor $v$ is a function of the ratio between the major and minor semiaxes, which is approximated by $\frac{L}{r}$, with the chain radius $r$ and the chain length $L$. $\frac{V_h N_A}{M_w}$ is the hydrated volume per unit mass, and can be approximated as $\frac{r^2 L N_A}{M_w}$. We thus have $\frac{\Delta D}{-\Delta f} \approx v\left(\frac{L}{r}\right)\frac{r^2 L N_A}{M_w}$. For HA, $r_{\mathrm{HA}} \approx 0.5$ nm and $\frac{L_{\mathrm{HA}}}{M_{\mathrm{W,\,HA}}} = \frac{L_{\mathrm{HA\ disaccharide}}}{M_{\mathrm{W,\,HA\ disaccharide}}} \approx \frac{1\ \mathrm{nm}}{400\ \mathrm{Da}}$, and for dsDNA, $r_{\mathrm{dsDNA}} \approx 1$ nm and $\frac{L_{\mathrm{dsDNA}}}{M_{\mathrm{W,\,dsDNA}}} = \frac{L_{\mathrm{dsDNA\ basepair}}}{M_{\mathrm{W,\,dsDNA\ basepair}}} \approx \frac{0.34\ \mathrm{nm}}{660\ \mathrm{Da}}$. Thus, one would expect the $\frac{\Delta D}{-\Delta f}$ vs $L$ data for HA and dsDNA to superpose when (i) the *x* axis for the DNA data is compressed by a factor of $\frac{r_{\mathrm{dsDNA}}}{r_{\mathrm{HA}}} \approx 2$, and (ii) the *y* axis for the DNA data is expanded by a factor of $\frac{r_{\mathrm{HA}}^2 L_{\mathrm{HA\ disaccharide}}}{M_{\mathrm{W,\,HA\ disaccharide}}} \frac{M_{\mathrm{W,\,dsDNA\ basepair}}}{r_{\mathrm{dsDNA}}^2 L_{\mathrm{dsDNA\ basepair}}} \approx \left(\frac{0.5\ \mathrm{nm}}{1\ \mathrm{nm}}\right)^2 \times \frac{1\ \mathrm{nm}}{0.34\ \mathrm{nm}} \times \frac{660\ \mathrm{Da}}{400\ \mathrm{Da}} \approx 1.21$. Indeed, the magnitude of the Δ*D*/-Δ*f* ratios for the re-scaled dsDNA data (*solid orange diamonds*) compares remarkably well to the HA standards (*black spheres*) in the limit of short chains where contour lengths are inferior to the persistence lengths of the polymer chains (~4 nm for HA, ~50 nm for dsDNA). This suggests that the theoretical approach proposed by Tsortos *et al.* makes accurate predictions for the shorter GAG oligosaccharides which can effectively be treated as rigid rods.
